# A corticostriatal learning mechanism linking excess striatal dopamine and auditory hallucinations

**DOI:** 10.1101/2025.03.18.643990

**Authors:** Kaushik Lakshminarasimhan, Justin Buck, Christoph Kellendonk, Guillermo Horga

## Abstract

Auditory hallucinations are linked to elevated striatal dopamine, but their underlying computational mechanisms have been obscured by regional heterogeneity in striatal dopamine signaling. To address this, we developed a normative circuit model in which corticostriatal plasticity in the ventral striatum is modulated by reward prediction errors to drive reinforcement learning while that in the sensory-dorsal striatum is modulated by sensory prediction errors derived from internal belief to drive self-supervised learning. We then validate the key predictions of this model using dopamine recordings across striatal regions in mice, as well as human behavior across various tasks including a new hybrid learning task. Finally, we find that changes in learning resulting from optogenetic stimulation of the sensory-dorsal striatum in mice and individual variability in hallucinations in humans are best explained by selectively enhancing dopamine levels in the model sensory-dorsal striatum. These findings identify plasticity mechanisms underlying biased learning of sensory expectations as a biologically plausible link between excess dopamine and hallucinations.

## 2 Introduction

Hallucinations are false percepts frequently experienced by people with psychotic disorders. Despite increasing insights into their cognitive mechanisms, our limited understanding of hallucination mechanisms at a biological circuit level precludes improvements to treatments which are often ineffective or poorly tolerated (Sommer et al., 2012). A promising venue to address this urgent need is formulating an end-to-end theory explaining how alterations in neural circuits drive changes in cognitive computations leading to hallucinatory experience i.e., a theory bridging the biology-experience gap (Fletcher & Frith, 2009).

A growing body of work has linked hallucinations to altered adaptation to sensory statistics (Friston, 2005; Adams et al., 2013; Corlett et al., 2019). Conceptualizing perception as an inferential process that combines learned sensory expectations with sensory evidence to form a percept, this work implicates altered learning or overweighting of sensory expectations in hallucinatory perception. In particular, in noisy sensory environments (e.g., signal detection tasks) psychotic individuals with hallucinations consistently show exaggerated biases toward learned expectations regardless of the true sensory evidence (Moseley et al., 2016; Powers et al., 2017; Cassidy et al., 2018; Corlett et al., 2019; Alderson-Day et al., 2022). While these insights represent substantial progress in our understanding of the cognitive-computational basis of hallucinations, existing theories still lack a detailed, biologically plausible implementation at the neural circuit level—a key requirement for a mechanistic explanation to be complete and actionable through biological interventions (X.-J. Wang & Krystal, 2014).

An established aspect of the neurobiology of hallucinations and other psychotic symptoms is elevated striatal dopamine (Laruelle et al., 1996; Abi-Dargham et al., 1998; Cassidy et al., 2018, 2019; McKetin et al., 2013), with pro-dopaminergic agents worsening and anti-dopaminergic drugs improving this symptom via striatal dopamine blockade (Kesby et al., 2018; Yun et al., 2023). A complete model of hallucinations must therefore incorporate dopamine and striatal circuits, including their increasingly well-documented heterogeneity across the striatum. This is particularly important given that dopamine elevation in psychosis predominates in dorsal striatum (Howes et al., 2009; Kegeles et al., 2010; Mizrahi et al., 2012; McCutcheon et al., 2018). Some previous theories indeed centered around the functional role of dopamine excess in psychosis (Stevens, 1973; Kapur, 2003; Fletcher & Frith, 2009; Heinz & Schlagenhauf, 2010; Maia & Frank, 2017), although knowledge about dopamine heterogeneity has since soared. Specifically, in contrast with the well-established role of ventral striatal dopamine in reward learning, dopamine in certain parts of the dorsal striatum (including the so-called auditory striatum) responds to sensory features irrespective of reward (Menegas et al., 2017, 2018; Schmack et al., 2021), consistent with distinct anatomical projections from sensory cortex to this “sensory dorsal striatum” (Chen et al., 2021) and its causal contribution to perception (Xiong et al., 2015; Guo et al., 2018; L. Wang et al., 2018; Chen et al., 2022). Anatomical evidence of specialized corticostriatal loops involving association sensory cortex and sensory dorsal striatum in primates similarly supports a role in perception and hallucinations (Middleton & Strick, 1996). A plausible mechanistic account of hallucinations should thus admit such anatomo-functional heterogeneity, specifying the involvement of distinct striatal modules in hallucination generation.

Here, we take a theory-driven approach to constructing a circuit-level computational model of hallucinations. Critically, we first develop a normative circuit model applying constraints from cross-species anatomical, physiological, and behavioral data. Our central hypothesis is that sensory-dorsal-striatal dopamine signals violations in sensory expectations (i.e., sensory prediction errors). This inspires the corollary hypothesis that excess dorsal-striatal dopamine in hallucinating individuals biases learning of sensory expectation which in turn drives false (hallucinated) percepts. We first strengthen the theoretical validity of the hypotheses by demonstrating that an optimized model learns to represent the nigrostriatal dopamine signal as a sensory prediction error. We then validate key predictions of the intact circuit model using dopamine recordings and optogenetic stimulation of the sensory striatum in behaving rodents, as well as human behavior. Finally, we simulate selective alterations in dopamine function and find that behavioral data in humans with varying degrees of hallucination propensity and clinical hallucination severity is consistent with model predictions of excess sensory dorsal-striatal dopamine function. Together, the results suggest that a learning- and corticostriatal-plasticity-based perspective of heterogeneous striatal dopamine signaling provides a unifying account of hallucination-related biological, cognitive, and phenomenological phenotypes across species.

## 3 Results

Clinical hallucinations manifest as an increased tendency to exhibit false alarms with high confidence in signal-detection tasks (Powers et al., 2017). These hallucination-related perceptual biases correlate with increased dorsal-striatal dopamine (Cassidy et al., 2018; Schmack et al., 2021). To investigate the mechanisms by which excess striatal dopamine may lead to high-confidence false alarms, we first take a computational view of the broader class of perceptual decision-making tasks. In such tasks, the state of the world is typically encoded in a series of noisy observations that unfold over time, and choosing an action yields a stochastic performance feedback or reward that depends on the state. One must therefore perform inference to estimate the underlying state from observations and then select an action based on the inferred state. The two processes impose unique constraints on learning. Optimal inference involves learning the statistics of the incoming stimulus (that is, *sensory expectation*) to minimize perceptual error, while selecting optimal actions in the inferred state requires learning the reward statistics (that is, *value*) to maximize reward (**Figure 1B**; Supplemental Figure 1). Since neither stimulus nor reward statistics are stationary in the real world, a normative approach to flexible perceptual decision-making should support learning of both types of statistics. Given that inference and action selection are cascaded processes, one could, in principle, bypass explicit learning of both the sensory expectation and value by relying on implicit trial-and-error learning based on performance feedback. However, previous studies have revealed engagement of both learning processes in tasks involving visual, temporal and auditory judgments, suggesting that perceptual decision-making is a two-stage process in which the first (sensory learning) stage can operate in the absence of external feedback while the second (reinforcement learning) stage requires feedback (Petrov et al., 2005; Dosher & Lu, 2017; Sohn & Jazayeri, 2021; Loewenstein et al., 2021). What biological mechanisms subserve the two learning objectives?

**Figure 1:**
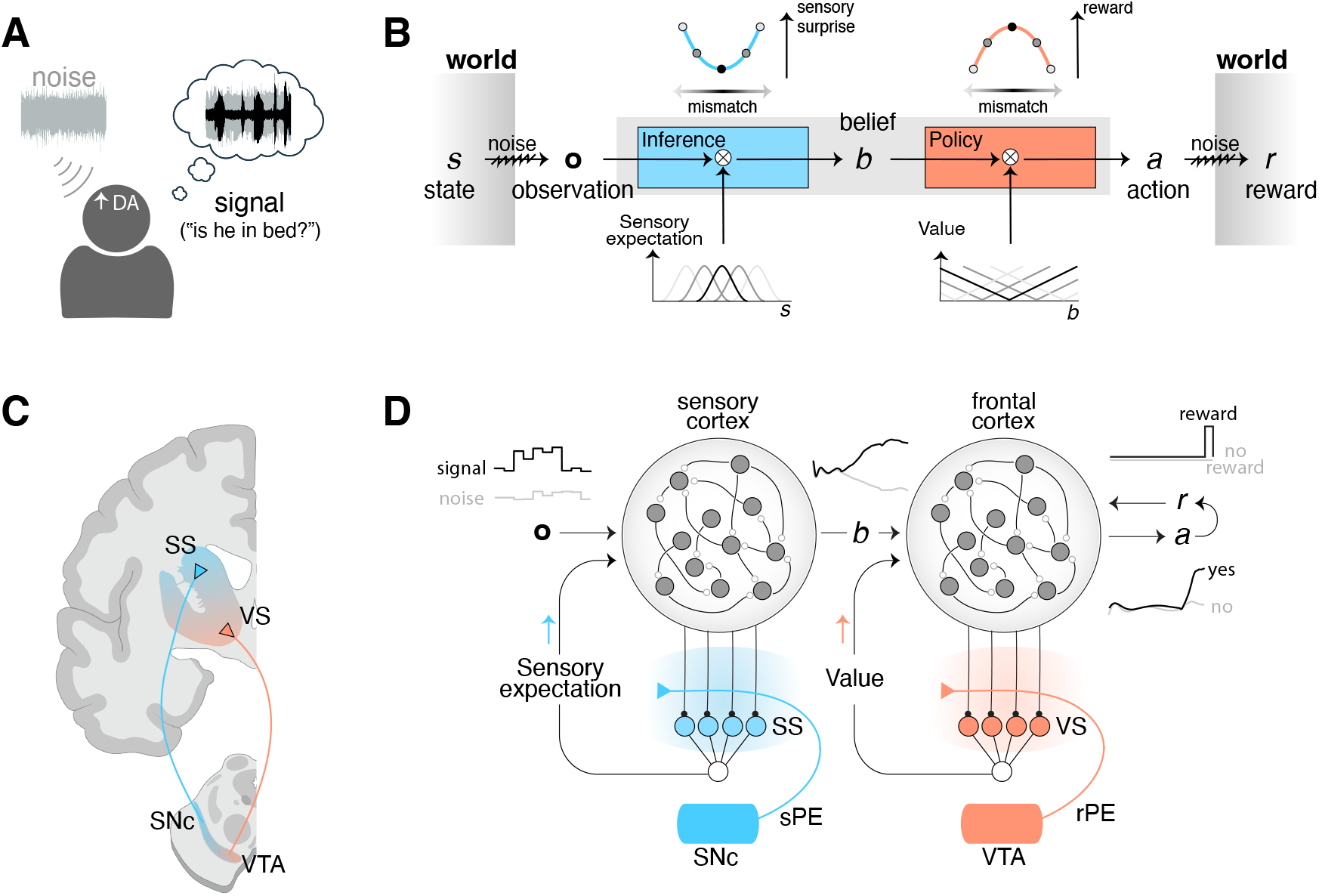
Modeling framework for auditory hallucinations. **A**. Excess striatal dopamine is linked to hallucinations, which typically consist of percepts with structured, specific content but consistently manifests as an increase in confident false alarms in signal detection tasks. **B**. Perceptual decision-making entails inference and action selection stages that must fulfill distinct desiderata – minimizing sensory surprise and maximizing reward, objectives that can be met by learning the sensory expectation and reward expectation (i.e., value) respectively. **C**. Distinct dopaminergic pathways target the ventral striatum and the sensory areas of the dorsal striatum. **D**. Circuit-level mechanisms of the computations in (B) can be implemented in a biologically plausible manner via plasticity mediated by distinct dopamergic prediction error signals in parallel corticostriatal loops. VS: ventral striatum, SS: sensory dorsal striatum, VTA: ventral tegmental area, SNc: Substantia nigra pars compacta, DA: dopamine, rPE: reward prediction error, sPE: sensory prediction error.

From a neurobiological standpoint, dopamine is known to support learning by modulating plasticity in corticostriatal synapses (Calabresi et al., 2007)(**Figure 1C** – left). However, existing models informed by reinforcement-learning theories focus primarily on the role of dopamine in value learning. In such models, learning is mediated by reward prediction errors (rPE) signalled by the mesolimbic pathway, i.e., midbrain dopamine neurons projecting from the ventral tegmental area to ventral striatum (VS). The role of dopamine signalling in the nigrostriatal pathway, i.e., from the substantia nigra to dorsal striatum is less clear. People with schizophrenia show elevated dopamine levels specifically in dorsal-striatal subregions targeted by this pathway, where dopamine synthesis and release capacity correlate with hallucination (and overall psychosis) severity (Laruelle et al., 1996; Howes et al., 2009; Kegeles et al., 2010; Cassidy et al., 2018, 2019). Furthermore, recent studies indicate dopamine responses to value-neutral stimuli in rodents (Takahashi et al., 2017; Stalnaker et al., 2019; Costa et al., 2025). Notably, neurons and dopamine signals in the nonmotor dorsal-striatal subregions play a causal role in perception (Guo et al., 2018; L. Wang et al., 2018; Chen et al., 2022), and dopamine stimulation in these subregions induces biased (hallucination-like) percepts (Schmack et al., 2021). For simplicity, we refer to this subregion as the sensory striatum (SS) throughout this work and use more specific terminology when referring to specific results from previous work. Based on the normative learning constraints outlined earlier, we hypothesized that rPE signals in VS dopamine facilitate learning of value while SS dopamine signals sensory prediction errors (sPE) – a specific type of prediction error appropriate for learning sensory expectation.

Temporal-difference reinforcement learning specifies the precise algebraic form of time-varying rPE signals but the composition of sPE signals remains ambiguous. One potential solution is provided by a class of models that rely on the discrepancy between actual and predicted state (Gläscher et al., 2010; Gardner et al., 2018), but constructing such prediction errors requires perfect knowledge of the stimulus identity which may not always be available to the animal. While rewards in instrumental tasks can be used to infer stimulus identity, this does not explain learning of stimulus statistics in the absence of feedback as noted earlier. Therefore, we asked whether sPE could instead be computed without external supervision. To address this, we derived an error signal for minimizing the cumulative change in the internal belief induced by sensory input, on average. Unlike feedback-dependent rPE signals which are useful for maximizing cumulative reward, the resulting sPE signals depended on the momentary changes in internal belief (Methods). To test whether this signal can mediate biologically plausible learning and to generate predictions at the neural level, we constructed a corticostriatal circuit model informed by neuroanatomical constraints (**Figure 1D** – right).

### 3.1 Modular corticostriatal model of perceptual decision-making

In the model, distinct midbrain nuclei—substantia nigra and ventral tegmental area—project to distinct striatal regions—SS and VS, respectively—to mediate plasticity in corticostriatal synapses according to a biologically plausible three-factor learning rule (Łukasz Kuśmierz et al., 2017; Gerstner et al., 2018). According to this rule, the update to a particular synaptic weight 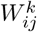 from cortical neuron *j* to striatal neuron *i* in region *k* ∈ {SS, VS} depends only on the pre- and post-synaptic activities, *r*_*j*_ and *r*_*i*_, and a prediction error signal *δ*_*k*_ conveyed by the midbrain dopamine inputs to that striatal region: 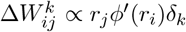, where *ϕ*(·) denotes neuronal nonlinearity (Methods). SS and VS neurons receive inputs from different cortical regions—sensory and frontal, respectively—and summed activities from striatal neurons in each region project back to the same cortical regions, forming parallel loops (Foster et al., 2021). While the precise anatomical pathway from the striatum to the cortex is complex, we use summed activity as a proxy since it has previously been shown to be a good indicator of the topographical organization of corticostriatal domains (Peters et al., 2021). Sensory and frontal cortices are modeled as recurrent neural networks, optimized respectively for perceptual inference and value-based decision-making (Methods).

Concretely, sensory observations are provided as external input to the model sensory cortex, which outputs a graded estimate of the posterior probability of the latent state i.e., belief to the model frontal cortex, which in turn outputs a binary yes/no response. Critically, both cortical computations depend on learning in the striatum. Accurate perceptual inference relies on learning of sensory expectations in the SS while accurate value-based decision depends on value learning in the VS. We assume this parallel learning is achieved by plasticity in the corticostriatal synapses gated by different types of dopamines signals in SS and VS, consistent with known differences in the sensory and frontal cortical projections to the midbrain dopaminergic neurons (Menegas et al., 2015; Sansalone et al., 2024). Concretely, plasticity in SS is modulated by the sPE signal derived above, where the momentary change in internal belief was calculated as the time derivative of the output of the sensory cortex (Methods). Plasticity in VS is modulated by a standard temporal-difference rPE signal following previous works (Rao, 2010; Hennig et al., 2023) (Methods).

To test the learning performance of this model, we varied stimulus or reward statistics in a signal-detection task by manipulating either the proportion of ‘signal’ trials or the fraction of rewarded correct ‘yes’ responses across blocks (**Figure 2A**; Methods). Although both manipulations disrupted performance initially following block transitions, the model adapted to them by adjusting its choices (**Figure 2B** – left). Critically, adaptation to manipulations of stimulus and reward statistics was mediated almost exclusively by changes in the output of SS (encoding sensory expectation) and VS (encoding value), respectively (**Figure 2B** – middle vs. right). This change of striatal outputs across trials was driven by changes in model SS dopamine (sPE) and VS dopamine (rPE), which gradually stabilized to baseline levels as learning converged (Supplemental Figure S2A). A similar dissociation was also observed within each trial, wherein stimulus and reward manipulations induced lasting changes in stimulus-evoked dopamine transients within the model SS and VS, respectively, albeit in opposite directions. Increasing the base rate of signal trials decreased the amplitude of signal-evoked SS dopamine after learning (**Figure 2C** – left), whereas increasing the reward probability associated with a correct ‘yes’ response increased signal-evoked VS dopamine (**Figure 2C** – right). This dichotomy in striatal dopamine was reflected in the model neural activity dynamics that emerged after learning. Although both types of manipulations produced comparable changes in choice-related response dynamics leading to similar levels of adaptation (stimulus → action; **Figure 2D** – left), manipulating stimulus statistics exclusively modified the initial state of the sensory cortex affecting belief computation in the sensory corticostriatal loop (stimulus → belief; **Figure 2D** – middle), while reward manipulation modified value dynamics that gate policy computation in the frontal corticostriatal loop (belief → action; **Figure 2D** – right).

**Figure 2:**
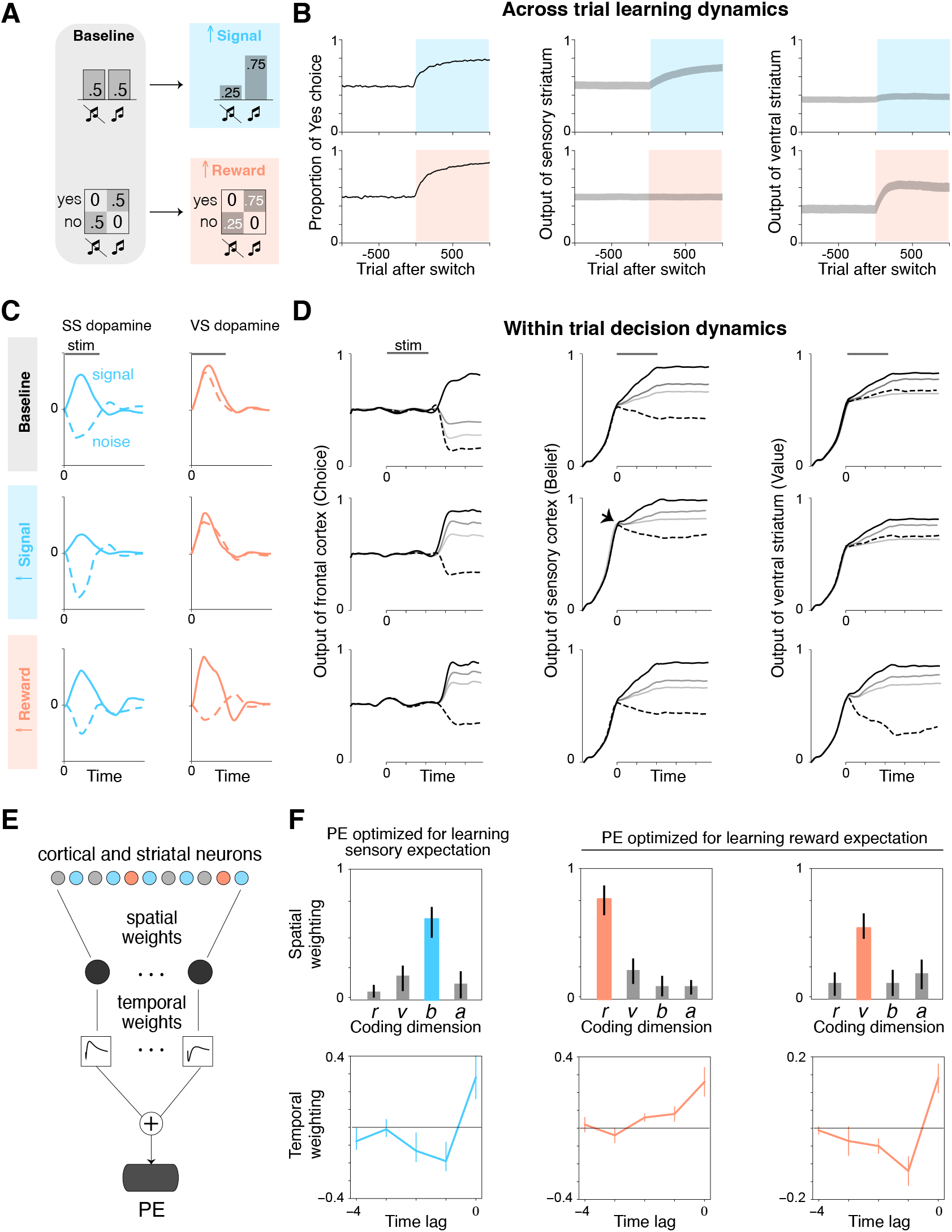
Distinct computational mechanisms of adaptation to signal and reward statistics. **A**. Starting from a block of baseline trials (signal/noise stimuli presented in equal proportion and correct yes/no rewarded in equal proportion), stimulus and reward statistics were separately manipulated by increasing the fraction of signal trials (top) and increasing the probability of rewarding a correct ‘yes’ response (bottom) respectively. **B**. Evolution of average reward (left), sensory expectation reflected in the output of model SS (middle), and the reward expectation reflected in the output of model VS (right) across trials before and after manipulation of stimulus (top row) and reward (bottom row) statistics. **C**. Stimulus-evoked model dopamine transients in signal (solid lines) and noise (dashed lines) trials. Rows correspond to different conditions and columns correspond to different striatal sub-regions. **D**. Average within-trial dynamics of choice (left), belief (middle), and value (right) under different conditions (rows). Dashed line corresponds to noise trials while lines with gray hues correspond to signal trials, with darker hues denoting a higher signal-to-noise ratio. **E**. Prediction error (PE) was optimized by expressing it as a sum of multiple spatiotemporal filters. Each spatial filter corresponds to a weighted sum of the activity of units in the cortical and striatal regions and each temporal filter corresponds to a weighted sum of the output of the spatial filter across a moving window. **F**. Left panel: PE optimized for learning sensory expectation – magnitude of alignment of the optimized spatial weights with activity dimension encoding reward (r), value (v), belief (b), and action (a), and the corresponding temporal weights. Right two panels: Similar to the left panel, but showing weights optimized for learning reward expectation. Error bars denote 95% confidence intervals obtained by bootstrapping across simulations.

Above results suggest that sPE constructed from momentary changes in internal belief can support a biologically plausible form of self-supervised learning of sensory expectations, alongside learning of value from temporal-difference rPE. We aimed to determine whether learning sensory expectations using the specific type of teaching signal assumed above, namely the time-derivative of internal belief, is normative. To address this, we expressed the prediction error encoded by striatal dopamine as an arbitrary spatiotemporal transformation of cortical and striatal activity and then used gradient-descent to optimize the spatial and temporal kernels separately to maximize either perceptual accuracy or reward (**Figure 2E**; Methods). Since this procedure seeks to optimize the teaching signal rather than task performance, it corresponds to meta-learning or learning-to-learn (Hospedales et al., 2022), a technique that has previously been applied to many neural systems (Wang et al., 2018; Jiang & Litwin-Kumar, 2021; Lakshminarasimhan et al., 2024). We found that a one-dimensional teaching signal was sufficient to optimize the model for perceptual accuracy (Supplemental Figure 2B – left). The resulting spatial weights were aligned primarily with the dimension of neural activity encoding the belief that the stimulus contained a signal, and the temporal weights resembled a derivative operator (**Figure 2F** – left; Methods). Alignment with dimensions that encoded reward, value (i.e., expected reward), and action was much weaker. This suggests that sPE constructed using momentary changes in internal belief optimizes biologically plausible learning of sensory expectations in this circuit architecture. In contrast, a similar approach to optimize the model for reward required a minimum of two spatiotemporal filters i.e., a two-dimensional teaching signal (Supplemental Figure 2B – right) that together constituted the temporal-difference rPE signal: one for selecting the current reward, and the other for extracting the time-derivative of value (**Figure 2F** – middle and right).

### 3.2 Signatures of sensory prediction errors in sensory striatal dopamine

We asked whether model VS and SS dopamine signals can inform the interpretation of dopamine signals recorded in those areas across different tasks. We considered two studies in which dopamine recordings were performed both in the SS (specifically, the rodent tail of striatum in this section) and VS under different conditions, allowing for a direct comparison of dopamine dynamics between the model and data.

In the first of these studies, (Menegas et al., 2017) tracked dopamine dynamics across the SS and VS during associative learning. They tested whether repeated training affected the observed pattern in SS and VS dopamine by introducing a new odor paired with a reward every day, and then measured dopamine activity while learning a new odor-reward association (**Figure 3A**; Methods). SS dopamine exhibited a response to novel odors which decreased across repeated exposures. In contrast, VS dopamine initially responded to rewards and only later on to odors in a manner that reflected the learned association between odors and rewards (**Figure 3B** – left). Simulating the model under associative learning yielded a strikingly similar pattern where odor-induced SS and VS dopamine dynamics decreased and increased respectively across trials (**Figure 3B** – right). The computational role of dopamine in the model offers an interpretation for these experimental results. Because model VS dopamine encodes rPE, the trajectory of dopamine signals reflects the backward progression of this temporal-difference error from rewards to odors in accordance with standard reinforcement learning theories. In contrast, SS dopamine encodes sPE and reflects the animal’s sensory surprise upon encountering an unexpected stimulus, which gradually reduces with repeated presentations as the animal comes to expect this stimulus. This qualitative finding was robust to the learning horizon determined by the discount factor (Supplemental Figure S3A). Note that we treat reward delivery itself as another sensory input in the above simulations (Methods) since the weak reward-related dopamine signals in SS have been attributed to the click of the water valve used to deliver reward (Menegas et al., 2018).

**Figure 3:**
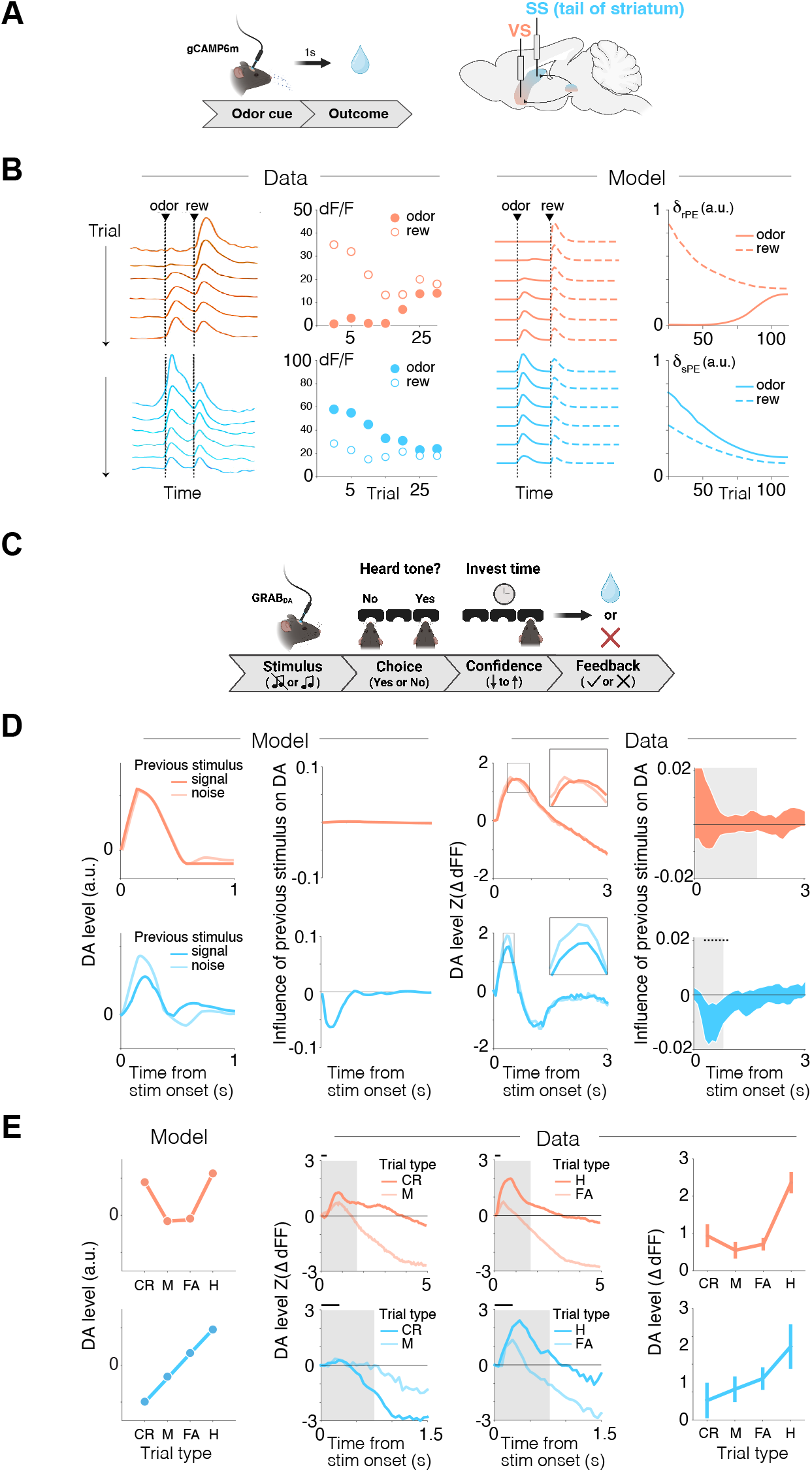
Signatures of distinct prediction errors in associative learning and signal detection. **A**. Schematic of the associative learning task. Mice were exposed to repeated pairings of a novel odor and reward delivery. **B. Left**: Dopamine dynamics in the mouse VS (top, orange, *n* = 9) and SS (bottom, cyan, *n* = 11) across trials. **Left middle**: Evolution of the amplitude of dopamine transients evoked by odor (open circles) and reward (filled circles). **Right**: Similar to the left panels but showing the evolution of rPE (top, orange) and sPE (bottom, cyan) in model simulations. **C**. Schematic of the signal detection task. Following stimulus presentation, mice reported their choice and confidence, estimated as the amount of time invested in the choice, and received feedback in the form of a reward. **D. Left**: Model-predicted stimulus-evoked rPE (top) and sPE (bottom), conditioned on the previous trial’s stimulus. **Left middle**: Regression weights for moving-window regression (Methods) quantifying the influence of previous trial’s stimulus on the prediction errors over time relative to stimulus onset. **Right**: Similar to the left panels but showing average dopamine levels in VS (top, orange) and SS (bottom, cyan) across mice (*n* = 6 for each region). Dopamine signals are expressed as *dF/F* and subtracted from pre-stimulus baseline (see Methods). Error bars denote 95% confidence intervals. Dots denote the time points in which the regression weight is significantly different from zero. **E. Left**: Time-averaged stimulus-evoked rPE (top, orange) and sPE (bottom, cyan) in the model simulations during trials classified based on stimulus-response relationship (CR: correct reject, M: miss, FA: False alarm, H: Hit). **Left middle**: Stimulus-evoked dopamine dynamics in VS (top) and SS (bottom) averaged across mice during CR trials and M trials. **Right middle**: Similar to the left middle panel, but during H and FA trials (*n* = 4 mice with recordings from both regions). **Right**: Time-averaged dopamine levels during the time windows denoted by the gray shaded regions in the middle panels, where window lengths are matched to the timescale of autocorrelation in dopamine signals in the two regions. Error bars denote standard error of the mean across mice and are normalized using the Cosineau method to remove between-subjects variance.

In a different study, (Schmack et al., 2021) recorded dopamine dynamics in the same striatal regions during auditory signal detection (**Figure 3C**). In this task, mice initiated trials with a center nosepoke, followed by a nosepoke at a left or right port depending on whether it heard a tone. To look for signatures of sPE and rPE in the data, we first examined dopamine signals in the model trained to perform signal detection. We found that the amplitude of signal-evoked model SS dopamine, but not VS dopamine, was modulated by the previous trial’s stimulus identity – in addition to scaling with signal strength (SNR), which has been previously documented (Schmack et al., 2021). Specifically, encountering a signal trial increases the expectation of a tone, decreasing the magnitude of signal-evoked response in SS dopamine (i.e., the sPE) in the subsequent trial (**Figure 3D** – left). We re-analyzed the data and found this precise pattern of modulation by stimulus history in the mouse SS, but not mouse VS, when accounting for the differences in the intrinsic timescale of dopamine dynamics (Supplemental Figure S3B) (**Figure 3D** – right): regression analyses showed that the previous trial’s stimulus identity (signal/noise) negatively modulated dopamine signal during the stimulus window in SS but not VS across all trials and mice (*s*_*t*−1_; VS: *β* = −0.076, 95% CI [−0.18, 0.022], *p* = 0.15; SS: *β* = −0.052, 95% CI [−0.087, −0.017], *p* = 0.0030). While the stimulus-history effect in SS dopamine is consistent with sPE signaling, we asked whether this signal specifically reflects fluctuations in subjective belief about the presence of a signal i.e., signed sensory surprise, as predicted by the model. When trials were grouped by stimulus-response relationship in the order of increasing confidence about the presence of a signal (CR *<* M *<* FA *<* H), the amplitude of stimulus-evoked SS dopamine increased monotonically. In contrast, the amplitude of model stimulus-evoked VS dopamine was higher on correct trials (CR and H) than incorrect trials, consistent with rPE signaling (**Figure 3E** – left). We found the same pattern in the subset of mice with recordings from both regions: stimulus-evoked dopamine increased monotonically across conditions ordered by confidence in the SS (condition, *x*: *β* = 0.21, 95% CI [0.010, 0.31], *p <* 5 *×* 10^−4^; *x*^2^ : *β* = 0.036, 95% CI [−0.080, 0.15], *p* = 0.52), and showed a U-shaped quadratic effect on condition in VS (*x*: *β* = 0.11, 95% CI [−0.0026, 0.21], *p* = 0.056; *x*^2^ : *β* = 0.30, 95% CI [0.18, 0.41], *p <* 5 *×* 10^−4^; *x*^2^ *×* region: *β* = 0.26, 95% CI [0.23, 0.29], *p <* 5 *×* 10^−4^; **Figure 3E** – right).

The above results show that dopamine dynamics in the mouse SS but not VS resembles model sPEs. We demonstrated earlier that this prediction error signal is optimized for learning sensory expectations via plasticity mechanisms. If this is the case, then perturbing SS dopamine should have a qualitatively similar effect on behavior as that on the model performance. This leads to two specific predictions. First, such perturbation should induce a bias in signal expectation rather than value. Second, the effect of this perturbation should gradually build up over time. We tested both predictions by re-analyzing the data from optogenetic stimulation experiments carried out during the same paradigm. Briefly, dopamine terminals in the SS were chronically stimulated via a laser across blocks of approximately fifty trials, interleaved with baseline blocks in which the laser was turned off (**Figure 4A**; Methods).

**Figure 4:**
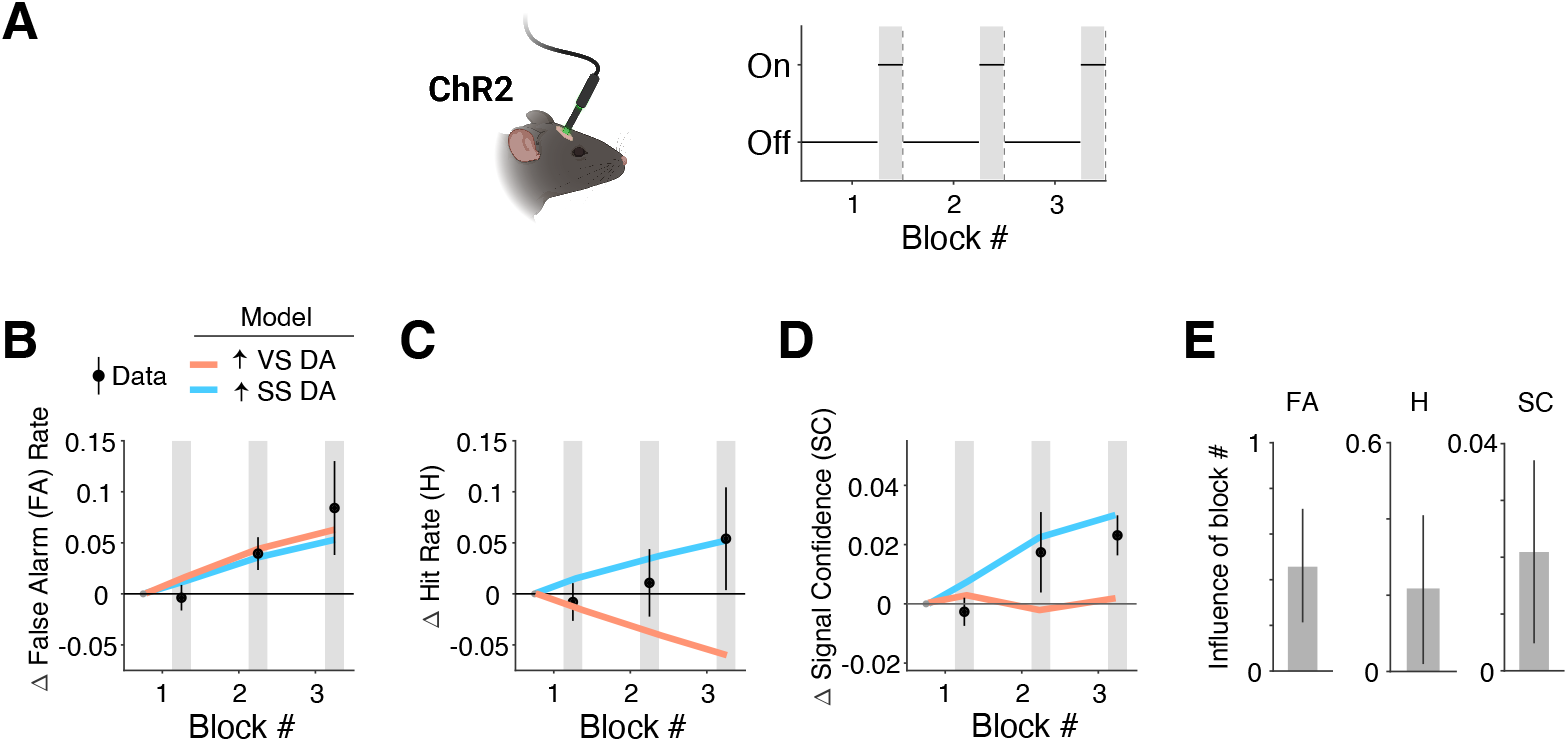
Evolution of signal expectation from SS dopamine stimulation. **A**. Schematic of the optogenetic stimulation protocol (*n* = 7). Laser was ON or OFF throughout the respective blocks. **B**. Change in the rate of false alarms with respect to the baseline (i.e., the first block of the experimental session) in each block. **C**. Similar to B, but showing the change in the rate of hits. **D**. Similar to B, but showing the change in signal confidence.**E**. Regression coefficient quantifying the effect of the block number on the false alarm rate (FA), hit rate (H), and signal confidence. For B, C, and D, error bars denote standard error of the mean across mice; for E, error bars denote 95% confidence intervals.

Model simulations indicated that VS dopamine stimulation increases the rate of false alarms and misses, whereas SS stimulation increases false alarms, hits, as well as the confidence that a signal is present, i.e., signal confidence (**Figure 4B-E** – orange vs cyan solid lines). This finding was robust to the precise form of perturbation applied to the dopamine prediction error signal (Methods; Supplemental Figure 4). In the mouse stimulation data, we found that, on average, stimulation significantly increased the rate of false alarms, as reported previously (Schmack et al., 2021) (Stimulation: *β* = 0.27, 95% CI [0.13, 0.39], *p <* 5 *×* 10^−4^). Moreover, stimulation also increased the signal confidence across all trials (Stimulation: *β* = 0.11, 95% CI [0.053, 0.17], *p <* 5*×*10^−4^), consistent with the hypothesis that SS dopamine stimulation influences behavior by increasing signal expectation rather than value. To test whether these effects were due to a learning rather than an instantaneous effect (driving role) of dopamine, we estimated how the above response measures evolved across blocks (47 *±* 9 trials per block). We found that false alarms, hits, signal confidence all exhibited a significant increase from baseline as a function of blocks within a stimulation session (**Figure 4B-E**; false alarms across session, *β* = 0.46, 95% CI [0.22, 0.71, *p <* 5 *×* 10^−4^; hits, *β* = 0.22, 95% CI [0.024, 0.41], *p* = 0.028; signal confidence, *β* = 0.021, 95% CI [0.0051, 0.037], *p* = 0.010). Notably, increasing model VS dopamine predicts an increase not only in the rate of false alarms but also misses, the latter of which is contradicted by data (**Figure 4C** – orange). Taken together, these analyses suggest that dopamine-mediated plasticity mechanisms in the SS play a role in learning sensory expectation.

### 3.3 Perceptual decision-making engages a two-stage learning process in humans

The results from the optogenetic stimulation study demonstrate distinct behavioral signatures of excess dopamine in the sensory striatal regions beyond changes in false alarm rates. Previous studies have noted that some of these signatures, such as signal confidence, correlate with hallucination propensity and clinical hallucinations in humans (Powers et al., 2017; Schmack et al., 2021). However, the extent to which hallucinatory experiences may be caused by an increase specifically in sPE signaling and whether they reflect an altered learning process rather than a constant perceptual bias remain unclear. This is in part because previous studies have typically investigated behavioral adaptation to either sensory or reward statistics but not both.

Motivated by the modular architecture in which stimulus and reward statistics are learned in distinct corticostriatal pathways, we developed a variant of a signal-detection task to identify unique signatures of learning sensory and reward expectations (**Figure 5A** – left). On each trial, participants experienced an ambiguous auditory stimulus (i.e., 1 kHz tone of varying SNR) and reported whether they heard a tone as well as the confidence associated with their judgement. Participants then received feedback about whether a tone was present and whether their choice resulted in a reward bonus. To engage both sensory and reward-based learning mechanisms, we manipulated both whether signals are more likely to be present or absent and whether a ‘yes’ choice is likely to produce feedback indicating more or less reward (**Figure 5A** – right).

**Figure 5:**
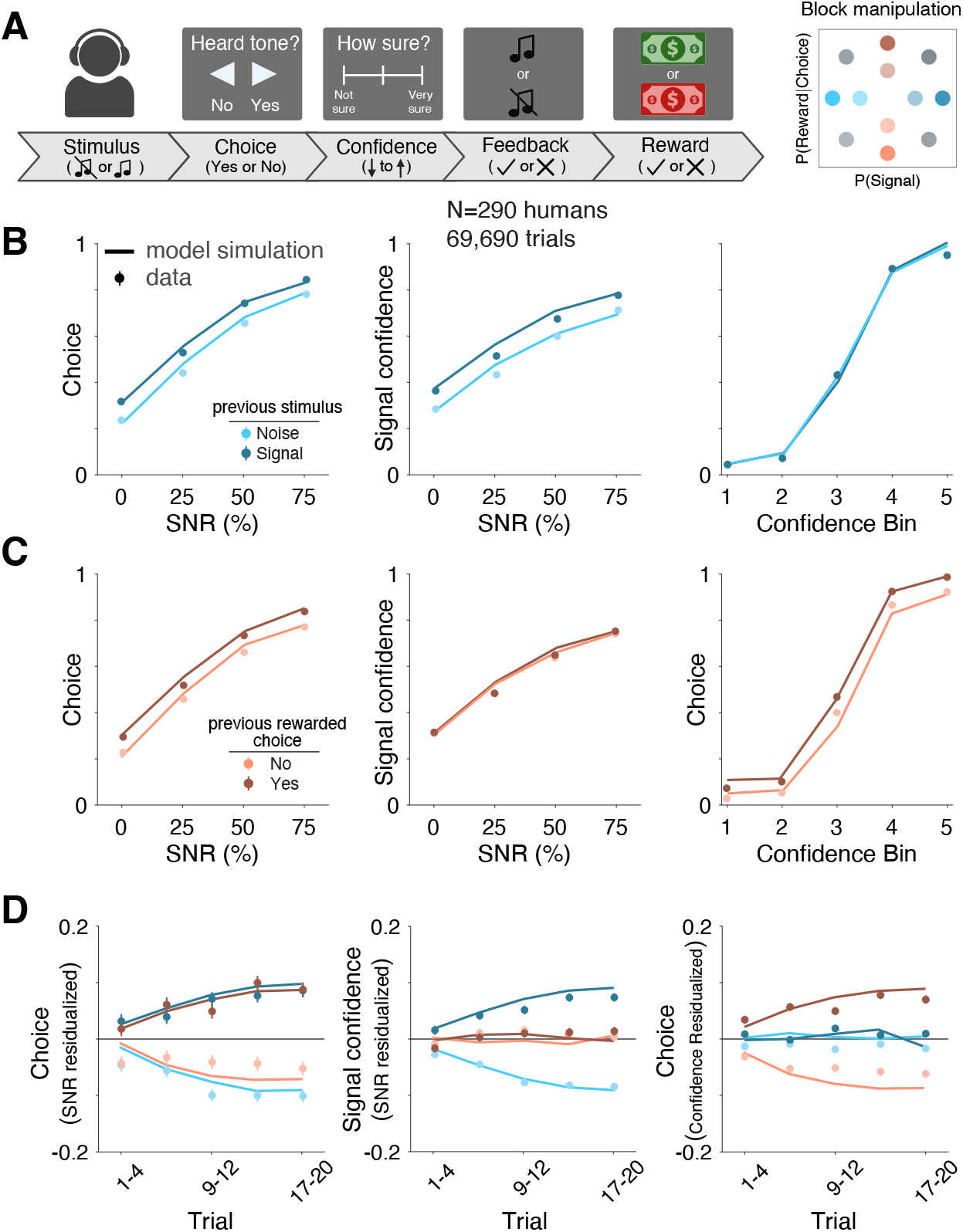
Distinct behavioral signatures of sensory and reward learning. **A**. Schematic of the task. **Left**: Human participants made a binary judgement about whether they heard a tone, reported the associated confidence on a continuous scale, and received feedback. **Right**: The fraction of signal trials and the fraction of rewarded yes responses were manipulated across blocks. **B**. Effect of signal history. **Left**: The fraction of yes choices as a function of the signal to noise ratio (SNR). **Middle**: Signal confidence as a function of SNR. **Right**: Participants’ policy, quantified as the fraction of yes choices as a function of confidence. **C**. Similar to B, but showing the effect of reward history. **D**. The evolution of choice, signal confidence, and policy across trials of the block for different manipulations. Error bars denote standard error of the mean across participants.

The model predicts that each of the two manipulations will drive distinct patterns of behavior in this task. As we show later, the manipulations gradually modify the pattern of behavioral responses across trials, implicating a slow process in which stimulus and reward statistics are learned in an incremental manner. Sensory and reward learning also manifest at a more granular level as distinct trial-history effects in the model. First, since both sensory and reward learning pathways influence choice via feedback (striatal output) to the cortex, the model predicts that choices would shift in accordance to the previous trial’s stimulus and reward (**Figure 5B,C** – left, solid lines). Indeed, human participants’ choices were biased both by the previous trial’s stimulus and previous trial’s reward (**Figure 5B,C** – left, filled circles; *s*_*t*−1_ : *β* = 0.41, 95% CI [0.38, 0.45], *p <* 5 *×* 10^−4^; *r*_*t*−1_ : *β* = 0.35, 95% CI [0.32, 0.39], *p <* 5 *×* 10^−4^). Second, since only the sensory-learning pathway (SS) influences belief estimates by conveying the sensory expectation, the model predicts that signal confidence would shift in alignment with the previous trial stimulus (**Figure 5B** – middle, solid lines) but not previous reward feedback (**Figure 5C** – middle, solid lines). Consistently, participants’ signal confidence was more strongly influenced by the previous stimulus than previous reward feedback (**Figure 5B,C** – middle, filled circles; Contrast Estimate = 0.22, 95% CI [0.20, 0.24], *p <* 5 *×* 10^−4^). Third, only the reward-learning pathway (VS) influences the policy, defined as the action taken in a given belief state and estimated as the fraction of yes choices binned by signal confidence for each participant. The model predicts that policy should shift in congruence with the previous reward feedback (**Figure 5C** – right, solid lines) but not previous stimulus (**Figure 5B** – right, solid lines). In line with this, participants’ policy was more strongly influenced by previous reward feedback than previous stimulus (**Figure 5B,C** – right, filled circles; Contrast Estimate = 0.85, 95% CI [0.76, 0.92], *p <* 5 *×* 10^−4^). Overall, this alignment with model predictions reveals dissociable signatures to changes in sensory and reward statistics.

In the model, adaptation to statistics is achieved by dopamine-mediated plasticity mechanisms. To further confirm that human participants engaged mechanisms that relied on a gradual learning process, we examined the evolution of all three variables—choice, confidence, and policy—across trials. We found that all quantities shifted gradually, plateauing towards the end of the block, suggesting that adaptations to each of the two manipulations is the outcome of slow learning processes (**Figure 5D** – left, P(Signal) *×* Trial: *β* = 0.066, 95% CI [0.047, 0.082], *p <* 5 *×* 10^−4^; P(Reward|Choice) *×* Trial: *β* = 0.069, 95% CI [0.053, 0.086], *p <* 5 *×* 10^−4^; **Figure 5D** – middle, P(Signal) *×* Trial: *β* = 0.039, 95% CI [0.032, 0.045], *p <* 5 *×* 10^−4^; P(Reward|Choice) *×* Trial: *β* = 0.0066, 95% CI [0.00041, 0.033], *p* = 0.033; **Figure 5D** – right, P(Signal) *×* Trial: *β* = −0.024, 95% CI [−0.050, 0.0032], *p* = 0.091; P(Reward|Choice) *×* Trial: *β* = 0.14, 95% CI [0.11, 0.16], *p <* 5 *×* 10^−4^). In addition to trial-specific choice and confidence reports, participants also periodically reported explicit subjective estimates of the underlying probability of signal and reward probability within each block of trials (Methods). In a high proportion of participants, estimates of both signal and reward probability were significantly modulated by the true block-level probabilities suggesting that they simultaneously learned both types of statistics (Supplemental Figure 5).

Lastly, since the model learns sensory expectations via self-supervision using its own evolving beliefs as a teaching signal, previous beliefs should influence the current choice even in the absence of feedback. To test whether humans engage in this type of learning, we re-analyzed data from two previous studies in which humans reported their decision confidence in addition to making binary judgments about stimuli without external supervision i.e., in the absence of performance feedback. In the first study, which used a signal detection paradigm (Powers et al., 2017), we found that confidence increased the propensity to repeat the choice made in the previous trial regardless of the signal-to-noise ratio of the stimulus in the current trial (**Figure 6A**; Confidence_*t*−1_: *β* = 0.51, 95% CI [0.42, 0.61], *p <* 5 *×* 10^−4^). We observed similar results in a separate study, which used a random dot motion paradigm to examine how participants learned the base rate of motion direction over trials in the absence of feedback (Zylberberg et al., 2018): choices made with high confidence increased the propensity to repeat that choice in the subsequent trial regardless of the coherence of the moving dots in that trial (**Figure 6B**; Confidence_*t*−1_: *β* = 0.64, 95% CI [0.56, 0.72], *p <* 5 *×* 10^−4^. The magnitude of the effect increased gradually with the level of confidence (Supplemental Figure 6A). These results suggest that humans employ a sensory learning mechanism guided by internal beliefs supported by dopamine-gated self-supervised plasticity.

**Figure 6:**
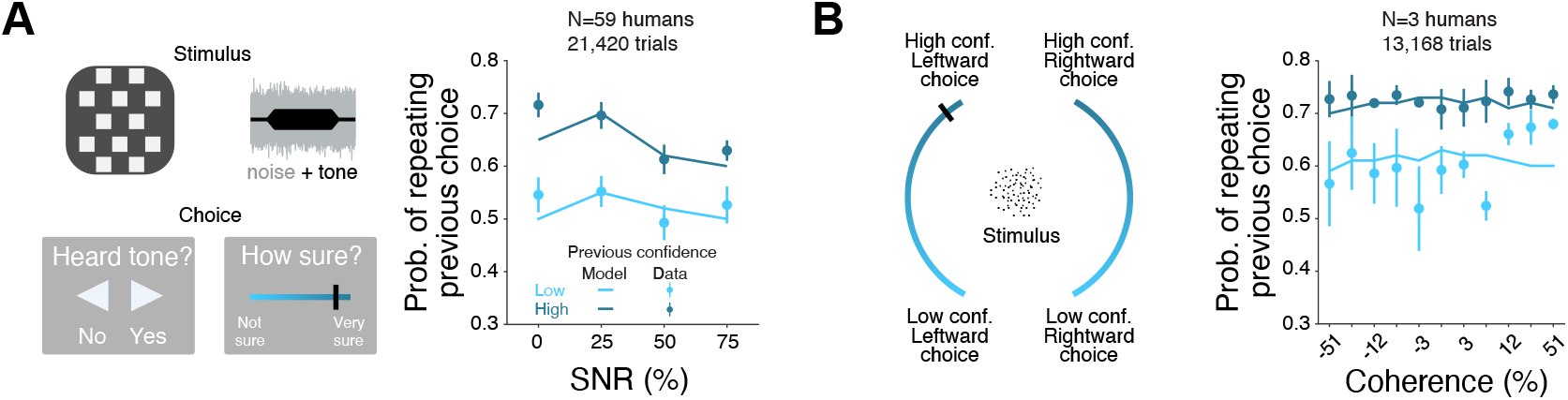
Behavioral signatures of self-supervised learning in the absence of feedback. **A**. Signal detection task. **Left**: Human participants made a binary judgment about whether they heard a tone during the presentation of a checkerboard pattern, reported the associated confidence on a continuous scale, and received no feedback. **Right**: The probability of repeating the previous trial’s choice (yes/no) as a function of current trial’s SNR, grouped by the confidence report in the previous trial (high/low). **B**. Motion discrimination task. **Left**: Participants made a binary judgment about the direction of moving dots displayed on a screen, reported the associated confidence on a continuous scale, and received no feedback. **Right**: The probability of repeating the previous trial’s choice (left/right) as a function of the coherence of the moving dots in the current trial, grouped by the confidence report in the previous trial (high/low). Solid lines correspond to model simulations and error bars denote standard error of the mean across participants.

### 3.4 Hallucination propensity is linked to increased sensory prediction errors

We now turn to the key question that motivated this study. What is the computational mechanism underlying the robust empirical link between hallucination severity and elevated striatal dopamine release? To address this, we combine insights from our analysis of the physiology of dopamine signaling in mouse striatum and history-dependent behavior in humans. Recall that stimulating dopamine terminals carrying signals that bear signatures of sPE in mice led to a slow build-up of signal confidence over the course of the experimental session. In humans, both stimulus and reward history biased future choices to a comparable extent but only the former also biased the confidence associated with the presence of a signal. Based on these findings, we reasoned that if hallucinations primarily result from increased sPE, then participants with increased hallucination proneness would also exhibit greater stimulus-history effects. We modeled the increase in sPE as a multiplicative perturbation of the prediction error signal (Methods) given work suggesting a disproportionate amplification of positive prediction errors in psychosis (Maia & Frank, 2017) (although results are robust to the precise form of perturbation).

To test this, using the data from human participants we asked whether the behavioral effects of stimulus history interacted with the participants’ hallucination proneness. In a sample (n=290) pre-screened to ensure a substantial proportion of individuals with high hallucination propensity (Methods), we confirmed the established relationship between higher hallucination propensity (CAPS) and increased false alarm rate (*ρ* = 0.16, *p* = 6.3 *×* 10^−3^). Leveraging our design to isolate stimulus-history effects, we found that hallucination propensity (CAPS) score was positively associated a tendency to report hearing a signal (including on noise trials, i.e., false alarms) following signal trials. (**Figure 7A,C** – left; *s*_*t*−1_ *×* CAPS: *β* = 0.074, 95% CI [0.037, 0.11], *p <* 5 *×* 10^−4^). Critically, this differential effect of stimulus history on choice was mirrored by a similar effect on signal confidence following signal trials (**Figure 7A,C** – middle; *s*_*t*−1_ *×* CAPS: *β* = 0.030, 95% CI [0.017, 0.044], *p <* 5 *×* 10^−4^), but not on the policy (**Figure 7A,C** – right; *s*_*t*−1_ *×* CAPS: *β* = 0.043, 95% CI [−0.0094,0.93], *p* = 0.11). These results indicating a selective bias in sensory learning with hallucination propensity are readily explained by elevated dopamine signaling specifically in the model SS (**Figure 7A** – solid lines), but not in the VS, suggesting that biased perception driving hallucination proneness could arise from elevated sPE in the SS. In contrast, we did not find a differential influence of reward history across hallucination propensity on choice (**Figure 7B,C** – left; *r*_*t*−1_ *×* CAPS: *β* = 0.0075, 95% CI [−0.026, 0.042], *p* = 0.65), signal confidence (**Figure 7B,C** – middle; *r*_*t*−1_ *×* CAPS: *β* = −0.0047, 95% CI [−0.017, 0.0082], *p* = 0.45), or the policy (**Figure 7B,C** – right; *r*_*t*−1_ *×* CAPS: *β* = 0.011, 95% CI [−0.043, 0.063], *p* = 0.64) and this was recapitulated by simulations in which dopamine in the model VS faithfully conveyed rPEs in both groups (**Figure 7B** – solid lines).

**Figure 7:**
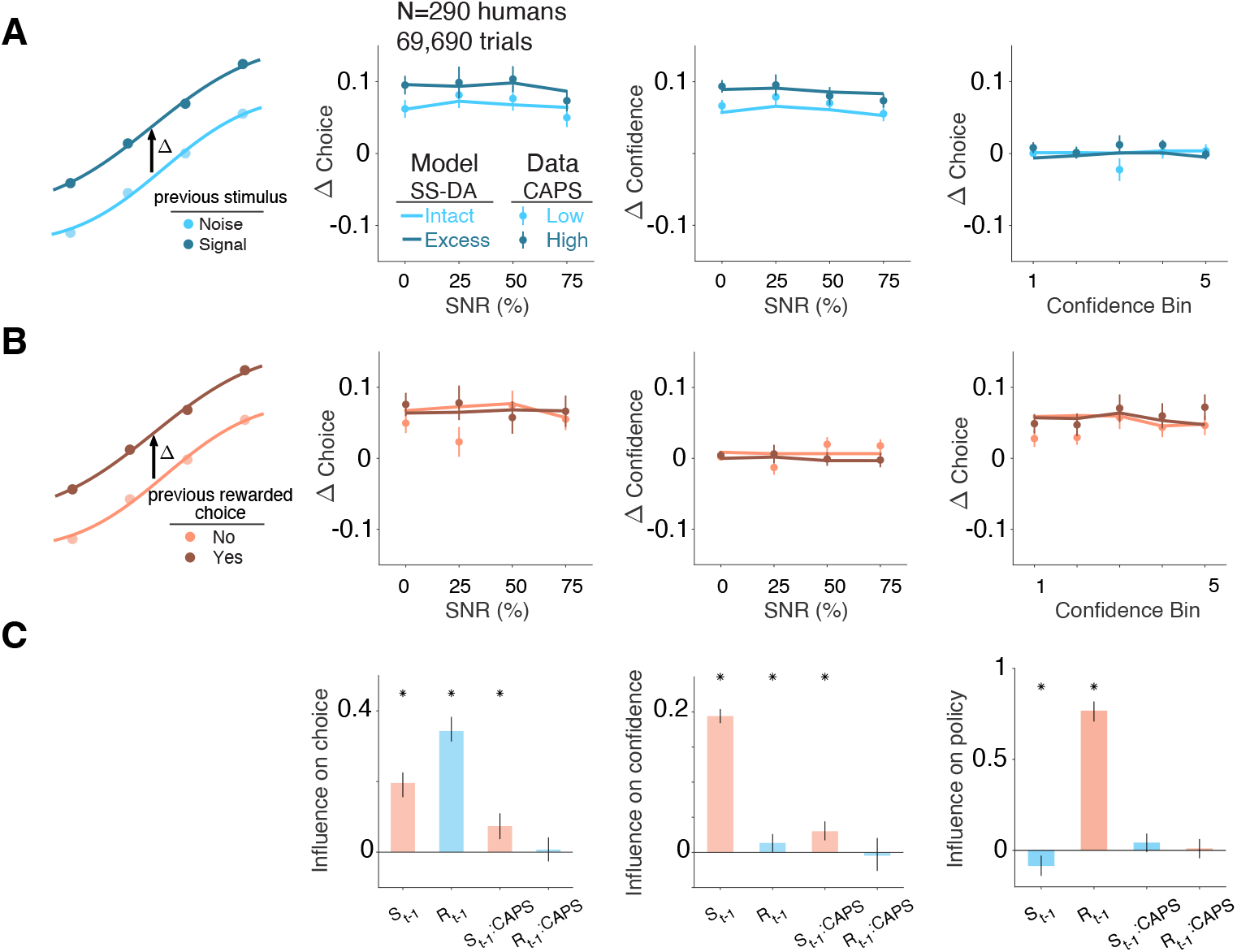
Exaggerated stimulus-history effects in humans with hallucination propensity. **A. Left**: The change in the rate at which participants report hearing a signal as a function of signal to noise ratio, after a signal trial, relative to after a noise trial. **Middle**: Similar to the left panel, but showing the change in the reported perceptual confidence. **Right**: Similar to the left panel, but showing the change in policy i.e., rate at which participants report hearing a signal as a function of confidence. Participants are grouped by their CAPS scores for visualization purposes (filled circles, Methods). Solid lines show model simulations in which sensory striatal dopamine was either intact (light) or increased (dark; Methods). **B**. Similar to A, but showing the change in response measures due to the previous trial’s reward feedback. **C**. Coefficients of a mixed effects model, showing the effect of previous trial’s stimulus (*s*_*t*−1_) and reward (*r*_*t*−1_), hallucination proneness (continuous CAPS score), and the interactions (*s*_*t*−1_ *×* CAPS, *r*_*t*−1_ *×* CAPS) on choice (left), confidence (middle) and policy (right). Asterisks denote *p <*0.05. For A and B, error bars denote standard error of the mean across participants and for C error bars denote 95% confidence intervals.

We asked whether altered sensory-history effects were specific to hallucination propensity rather than a reflection of general cognitive ability. To address this, we examined how trial-history effects interacted with a measure of participants’ fluid intelligence (Raven’s progressive matrices, Methods). Strikingly, we found an opposite pattern of results whereby reward history but not stimulus history was negatively associated Raven’s scores (Supplemental Figure 7A-C). The double dissociation between the effects of hallucination propensity and cognitive ability was further evident in a combined mixed-effects analysis which revealed that stimulus-history effects interacted with hallucination propensity (CAPS) and reward-history effects interacted with fluid intelligence (Raven’s) (Supplemental Figure 7D). Altogether, these results suggest that biased hallucination-like percepts are consistent with selective alterations in sensory learning and SS corticostriatal plasticity due to elevated sPE signaling.

These results raise the question of whether a similar mechanism is at play in hallucinating psychotic individuals diagnosed with schizophrenia, a population with demonstrated increase in dorsal-striatal dopamine function (Howes et al., 2009; Kegeles et al., 2010; Mizrahi et al., 2012; McCutcheon et al., 2018). To test whether sPE-based mechanisms can account for clinical hallucinations, we re-analyzed data from a previous study in which patients with schizophrenia with different levels of clinical hallucination severity performed an auditory conditioning task (Powers et al., 2017) (**Figure 8A**). Because patients on this task did not receive feedback, these data also allowed us to test the hypothesized hallucination-related bias in self-supervised sensory learning, which should emerge regardless of feedback. We found that modeling hallucinations as an increase in sPE was indeed sufficient to account for a broad range of behavioral differences across patients grouped by hallucination severity, including the previously reported increases in false alarms (conditioned hallucinations) and false-alarm confidence in patients with high hallucination severity. In addition to increased false alarms, the model captured an increase in hits observed in those with more severe hallucinations (**Figure 8B**; Severity: *β* = 2.00, 95% CI [0.98, 2.99], *p <* 5 *×* 10^−4^). Second, increasing the model’s sPE affected confidence in a choice-dependent manner, capturing an interaction between choice and hallucination severity in patients (**Figure 8C**; Severity *×* Choice: *β* = 0.15, 95% CI [0.058, 0.24], *p <* 5 *×* 10^−4^). Under our model, biased sensory expectations driving these effects result from alterations in sensory learning due to excess dopamine. Consistent with this prediction, we found a significant effect of block number on the rate of false alarms and signal confidence across both groups of patients (**Figure 8D**; Block number: *β* = 0.26, 95% CI [0.081, 0.43], *p* = 0.009), with qualitative differences between groups explained by solely increasing model sPE. The hallucination-severity effects in patients were also recapitulated by a model where prior weighting increased with hallucination severity (Supplemental Figure 8; Methods) suggesting that findings from the proposed corticostriatal learning model align with previous accounts of hallucinations in terms of overweighting of sensory expectations (Powers et al., 2017). These results thus provide a dopaminergic and corticostriatal-plasticity basis for prior overweighting, bridging circuit-level and algorithmic accounts of clinical hallucinations.

**Figure 8:**
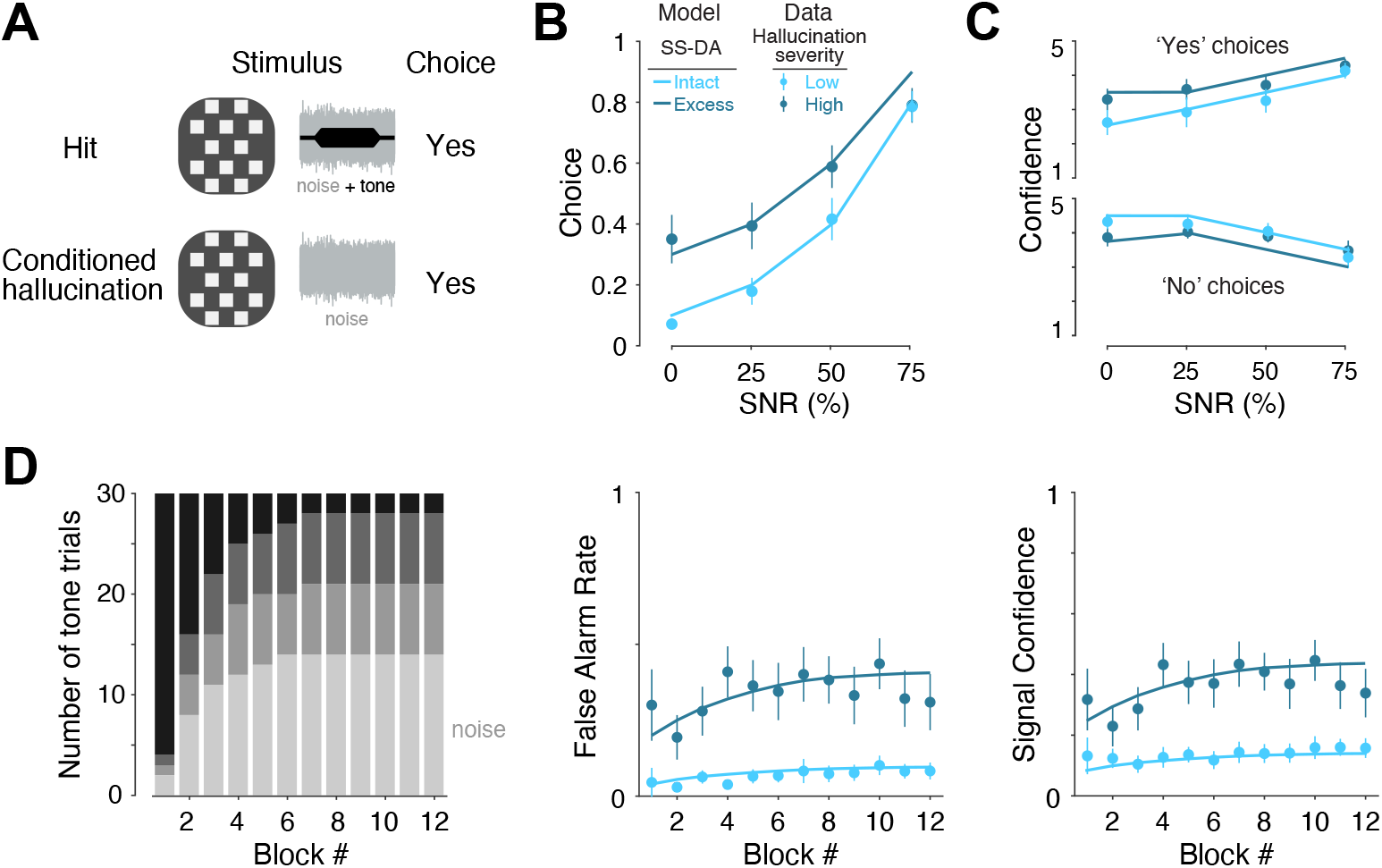
Increased sensory prediction error recapitulates behavior of hallucinating patients. **A**. Human participants made a binary judgment about whether they heard a tone during the presentation of a checkerboard pattern, reported the associated confidence on a continuous scale, and received no feedback. **B**. The change in the rate at which patients with low (*n* = 14) and high (*n* = 15) hallucination severity report hearing a signal as a function of signal to noise ratio. **C**. Similar to **B**, but showing the change in the reported perceptual confidence separately for trials in which participants choose ‘Yes’ (top) and ‘No’ (bottom). **D**. The number of noise and tone trials (left), the rate of false alarms (“conditioned hallucinations”, middle), and the average signal confidence (right), as a function of the block number. Patients are grouped by their hallucination severity (light vs. dark, Methods). Solid lines show model simulations in which sensory striatal dopamine was either intact (light) or increased (dark; Methods). Error bars denote standard error of the mean across participants.

While the model explains a link between increased SS dopamine and hallucinations in settings with binary states, hallucinatory experiences in patients with psychosis typically have complex structure: they are not just a mistaken belief that a signal is present but a belief with specific perceptual content (e.g., hearing a concrete phrase; **Figure 1A**). While a detailed model of complex auditory processing is beyond the scope of this work, we asked whether the model could explain biased beliefs about stimulus categories in a multi-class discrimination setting. To test this, we developed a model variant to perform a ten-way classification task in which each class is defined by a predominant (high-amplitude) stimulus feature, assuming that sensory expectations for different classes are learned in parallel via feature-specific sPEs (**Figure 9A**), analogous to feature-specific rPE models (Lee et al., 2024). We found that a global increase in sPE resulted in heightened expectations specifically for categories that were relatively more common to begin with (**Figure 9B**). Consequently, the model with increased sPE developed a strong tendency to report the presence of the most common category (category 1) when presented with a blank stimulus i.e., false alarms, and the associated belief strength was more often comparable to that experienced during the presence of stimuli from one of the categories i.e., hits (**Figure 9C** – left). We quantified this overlap as 1 − *d*′ where *d*′ was estimated as the area under the receiver operating characteristic curve (Methods) and found that the model with excess sPE exhibited a substantially greater overlap between category-1 false alarms and hit trials than the model with intact sPE (**Figure 9C** – right). Furthermore, the content-specificity of the belief on individual trials, defined as the difference between the category-1 belief and the average belief for all other categories, was also substantially greater in the model with excess sPE (**Figure 9D**). Together, these results suggest that the computational logic that links excess striatal dopamine to hallucination-like percepts in simple signal detection settings can also capture biased beliefs toward specific contents seen in typical complex hallucinations.

**Figure 9:**
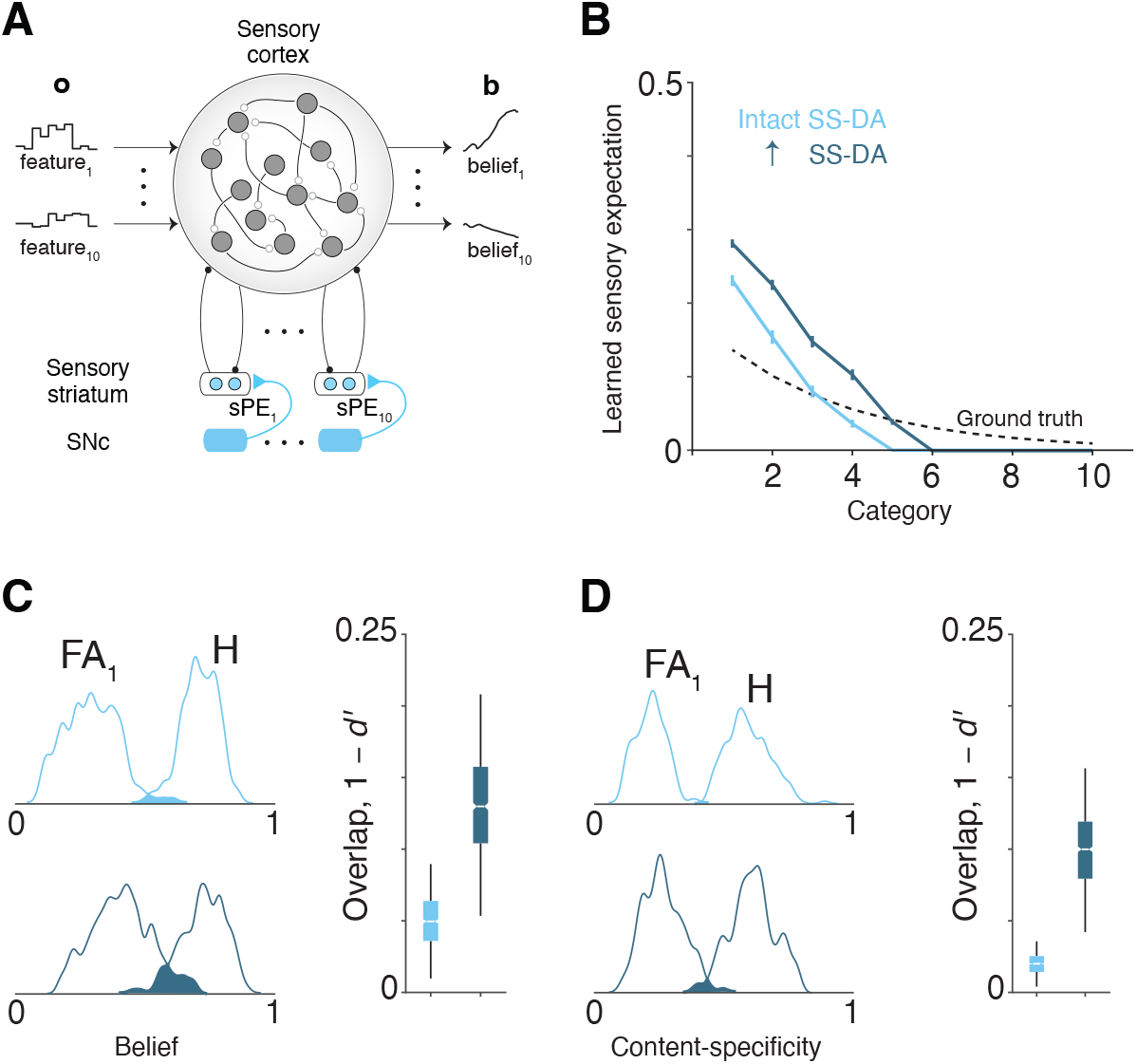
Increased sensory prediction error recapitulates content-specific hallucinations in multiclass classification. **A**. The model architecture modified to separately learn sensory expectations for each category using feature-specific sensory prediction errors (sPE). For simplicity, the action-selection module is not explicitly modeled here. **B**. The true category distribution (black) and the expectations learned by the model with intact (light blue) and excess (dark blue) sPE. **C**. Left: The distribution of posterior beliefs favoring the most common category (category 1) on blank trials (FA_1_) and that of posterior beliefs consistent with the stimulus category on non-blank trials (H) for the model with intact sPE (top) and excess sPE (bottom). Right: The overlap between the two distributions, quantified as 1 − *d*′ where *d*′ corresponds to the AUC (Methods). **D**. Similar to **D**. but showing the distribution of content-specificity, defined as the belief about the favored category relative to other categories (Methods). Error bars denote standard error of the mean across participants.

## 4 Discussion

We developed an anatomically constrained, normative circuit model of corticostriatal loops in which dopaminergic prediction-error signals in the ventral and dorsal sensory striatum are optimized to respectively support value learning and sensory learning via a plausible cortico-striatal-plasticity rule. Dopamine dynamics in the mouse dorsal sensory (tail) striatum mirrored sensory prediction errors essential for learning sensory expectation, and dopamine stimulation in this region gradually biased perceptual beliefs indicating a plasticity-based mechanism. In an auditory detection task, human behavior showed both sensory and reward learning, but only changes in the former correlated with hallucination proneness and additionally captured behaviors associated with clinical hallucinations. Thus, a single perturbation in a biologically constrained circuit model – enhanced dopaminergic sensory prediction errors in dorsal sensory striatum – is sufficient to explain hallucinations through a local change in cortico-striatal plasticity that biases internal beliefs about sensory states. These findings reveal a potential computational circuit mechanism underpinning the link between excess striatal dopamine and auditory hallucinations, and provide a unifying account of influential hallucination-related results across paradigms and species (Powers et al., 2017; Schmack et al., 2021; Corlett et al., 2019). In doing so, our model addresses a crucial gap in the development of computational circuit-level models of altered subjective experiences characteristic of psychiatric illness – here, the emergence of biased internal beliefs driving hallucinatory percepts (Fletcher & Frith, 2009; X.-J. Wang & Krystal, 2014).

Although the complex pathophysiology of psychosis points to a dysfunction in a broad range of synaptic mechanisms involving dopamine, glutamate, GABA, etc., almost all effective antipsychotic medications target striatal dopamine receptors, and dopamine stimulants induce psychotic symptoms in healthy individuals (Curran et al., 2004; Sommer et al., 2012; Lally & MacCabe, 2015). Unsurprisingly, some of the earliest theories of psychosis focused on dopamine (Stevens, 1973; Seeman & Kapur, 2000; Heinz, 2002; Kapur, 2003). Reflecting the dominant computational views at the time, these theories appealed to the role of dopamine in reward learning and motivational salience. However, the problems with these early theories were twofold. First, they did not directly model how excess dopamine leads to psychotic symptoms like hallucinations, instead relying on paradigms like associative and motor learning for empirical validation (Walter et al., 2009; Gradin et al., 2011). Second, they were not neurobiologically grounded in a circuit model and therefore carried limited mechanistic and translational relevance—with few notable exceptions (Maia & Frank, 2017). Consequently and despite established convergent support, state-of-the-art theoretical frameworks focused on pathological inference like those based on predictive processing have often sidelined the ‘dopamine hypothesis’ in favor of models of cortical imbalance between excitation and inhibition (Kehrer, 2008; Jardri & Denève, 2013; Sterzer et al., 2018; Keller & Sterzer, 2024). However, an emphasis on inferential mechanisms does not preclude a role for dopamine (Fletcher & Frith, 2009; Horga & Abi-Dargham, 2019). In fact, a growing body of basic research using modern molecular tools points to anatomically segregated and physiologically distinct dopamine signaling pathways, including a role in perception for the dorsal sensory striatum and dopamine therein (L. Wang et al., 2018; Guo et al., 2018; Schmack et al., 2021; Chen et al., 2022). This development has paralleled proposals that extend the canonical, reward-centric computational theories by rethinking dopamine signaling as a generalized prediction error (Gardner et al., 2018). Informed by these empirical and theoretical results, we developed a corticostriatal model in which dopamine contributes to the inferential process by enabling self-supervised learning of sensory expectations in the sensory striatum.

### 4.1 Novelty and implications

The proposed model of the sensory striatum differs from alternative models employing different types of outcome-specific prediction errors (Gardner et al., 2018; Lee et al., 2024) in that it allows for learning exclusively from internal beliefs, i.e., in the absence of actions and/or outcomes. This is in line with empirical work showing that humans (and other animals) can learn stimulus statistics in the absence of feedback (Petrov et al., 2006; Zylberberg et al., 2018; Fleming et al., 2019; Loewenstein et al., 2021; Schmid et al., 2024). A previous algorithmic account proposed that in the absence of feedback, sensory expectations must be updated on the basis of counterfactual beliefs evaluated using uniform priors to avoid self-reinforcement or double counting (Zylberberg et al., 2018). While a uniform prior is a natural choice for binary decision-making tasks in which the states e.g., motion directions are a priori equally likely, it is less intuitive when learning expectations for rare sensory stimuli. Furthermore, computing counterfactual estimates of belief places additional computational burden on the learning system. Learning via sPE also avoids belief self-reinforcement and offers a biologically plausible alternative for self-supervised learning without invoking the computation of counterfactuals (Supplemental Figure 6B).

Another advantage of sPE-gated corticostriatal plasticity is that it enables post-synaptic neurons, i.e., neurons in the sensory striatum to develop action-independent sensory representations, allowing for flexible reuse of these signals in different contexts. Action-independent coding has indeed been reported recently in the sensory striatum of rodents (Guo et al., 2018) and is consistent with earlier studies indicating that optogenetic stimulation of neurons in this region biases perception rather than choice (L. Wang et al., 2018). Because striatal learning influences cortical belief representations in the model, the proposed mechanism also explains history- and context-dependent reorganization of belief signals in the associative cortex (Akrami et al., 2018; Noel et al., 2022; Lakshminarasimhan et al., 2023). While the model postulates sPE-based learning in sensory neurons in the striatum in general, we validated it using data from the rodent tail of striatum. Previous studies demonstrated that dopamine signal in this region is consistent with other types of prediction errors such as action prediction errors (Miller et al., 2019; Greenstreet et al., 2022; Lakshminarasimhan, 2024) and threat prediction errors (Menegas et al., 2018; Akiti et al., 2022). We note that these proposals are not mutually exclusive as they pertain to different task dimensions – sensory input, motor command, and outcome valence – and dopamine dynamics could multiplex these signals to enable learning along different dimensions. Functional heterogeneity within subregions of the tail striatum is also possible.

Additionally, recent works have revealed that dopamine responses in the ventral striatum are influenced by the internal model of the environment (Daw et al., 2011; Starkweather et al., 2017; Babayan et al., 2018; Krausz et al., 2023). But the extent to which dopamine responses underlie the learning of those internal models remains unclear. The proposed model suggests a plausible mechanism by which model-free, self-supervised updates in the sensory striatum facilitates learning of the internal model, which subsequently influences dopamine signaling for reward-based learning. This parallel learning scheme can also explain the paradoxical finding that perceptual decision confidence and stimulus strength exert opposite effects on serial dependency in choices (Braun et al., 2018; Bosch et al., 2020; Lak et al., 2020). Whereas in the absence of feedback, large sPEs associated with high confidence choices strengthen the magnitude of serial dependency (**Figure 6**), correct judgments of ambiguous stimuli strengthen serial dependency by inducing large rPEs following unexpected reward feedback. Based on the segregation of sPE-based and rPE-based learning to the sensory and ventral striatum in the model, we propose that the previously reported link between association cortex–dorsal striatum connectivity and history effects reflects a specific role for this circuit in self-supervised learning (Hwang et al., 2019; Urai & Donner, 2022).

Our modeling approach represents an integration of normative and mechanistic considerations and is particularly appealing from the perspective of preclinical translational research since it allows for testing both granular predictions about neural dynamics and behavioral predictions using data across species. The proposed link between dopamine and sensory learning has major implications for understanding the neural basis of auditory hallucinations. First, we identified signatures of sensory prediction errors in the sensory striatum but not in the ventral striatum, a result that paralleled the selective interaction between hallucination proneness and stimulus history but not reward history in humans. A notable advantage of our new task design is that, improving upon previous designs, it allowed us to isolate sensory learning from response bias and general learning changes and specifically link sensory learning biases consistent with increased sPE to hallucination proneness. This selective change in sensory learning reconciles findings from PET imaging studies that localize elevated extracellular dopamine and synthesis capacity to the human dorsal striatum (Howes et al., 2009; Kegeles et al., 2010; Mizrahi et al., 2012) rather than ventral (limbic) striatum as anticipated by earlier theories of dopamine and psychosis (Kapur, 2003). Second, a specialized role for dopamine in this subregion vis-a-vis the non-region-specific antidopaminergic (D2-receptor blockade) effect of most antipsychotic drugs provides a potential explanation of their wide-ranging side effects in terms of off-target effects (Huhn et al., 2019). Lastly, our analysis of the time course of evolution of perceptual bias in response to dopamine stimulation was consistent with a plasticity-based mechanism. A role for sensory-striatal dopamine in sensory learning via a plasticity mechanism (rather than an instantaneous driving effect of dopamine on decisions) could potentially explain the slow clinical response to antipsychotic medications (Maia & Frank, 2017; Cassidy et al., 2018) and provides a biological basis for the development of strong priors in Bayesian-inference models of hallucinations (Corlett et al., 2019; Duhamel et al., 2023). More generally, a key insight from our work is highlighting the relevance of SS corticostriatal plasticity as a more proximal driver of psychotic symptoms (hallucinations), with increased SS dopamine being a mediator of plasticity changes (which also depend on glutamatergic cortical presynaptic input and GABAergic striatal post-synaptic output captured by the three-factor plasticity rule in our model). This suggests that pharmacological agents directly targeting corticostriatal plasticity (via dopaminergic or non-dopaminergic pathways) could be explored as a potential treatment for psychosis. Understanding the interaction between dopaminergic, glutamatergic, and cholinergic systems in corticostriatal synapses may provide a way to target this plasticity (Reynolds et al., 2022; Gallo et al., 2022; Chantranupong et al., 2023), which could also explain the efficacy of an emerging class of antipsychotic drugs based on muscarinic agonism (Mirza et al., 2003; Kaul et al., 2024).

SS dopamine may contribute to hallucinatory percepts through plasticity-based mechanisms while also playing a more direct role in modulating sensory expectations, as suggested by previous studies (Schmack et al., 2021). However, an exclusive role in encoding sensory expectations cannot account for the modulation of stimulus-evoked dopamine by stimulus history or the gradual impact of dopamine stimulation noted here (Figures 3D and 4). Under certain assumptions, in contrast, our model can also explain previous findings showing elevated pre-stimulus dopamine from an sPE perspective (Supplemental Fig. S3C). Likewise, while threat prediction errors and action prediction errors can gradually bias choices, they do not readily explain the characteristic perceptual qualities of hallucinations (e.g., why people with psychosis *hear* specific, well-defined sounds and voices) or the strong conviction of the associated beliefs (and control analyses speak against these alternative explanations for the data we analyzed here; Supplemental Fig. S3D). The proposed role for sensory striatal dopamine in learning sensory expectations thus provides a parsimonious link between excess dopamine and hallucinations.

### 4.2 Predictions and extensions

Our analyses of striatal dopamine revealed signatures of sensory prediction errors in the tail of the rodent striatum, but the model makes concrete predictions linking dopamine to the learning of sensory expectations that could be validated using signal-detection paradigms with changing signal probability. First, stimulus-induced SS dopamine levels should decrease as signal probability increases, since higher probability makes signals less surprising. Second, this decrease in dopamine should be accompanied by an increase in cortical activity within the dimension that encodes belief, driven by input from the basal ganglia prior to stimulus onset. Third, discrepancies between signal probability decoded from the initial cortical state and the true signal probability should correlate with deviations of the sensory striatal dopamine from the optimal sensory prediction error signal.

One challenge in testing the model predictions is the need to precisely target recordings to the corticostriatal circuit involved in learning and inference of the variable of interest. While in rodents the loop involving the primary auditory cortex is an appropriate choice in simple auditory signal-detection tasks, future extensions of the model should expand this approach to learn complex tasks using architectures that incorporate hierarchical processing of rich sensory inputs, informed by detailed wiring diagrams of the cortico-basal-ganglia loop (Foster et al., 2021). This will prove crucial to testing whether the pathophysiology of complex hallucinations in patients with psychosis is limited to dopamine signals that encode prediction errors about specific sensory features or if extends more broadly to semantic content. A related assumption was our treatment of dopamine as a scalar prediction-error signal. While this is justified and supported by empirical work in the reward domain, sensory inputs inherently comprise multiple feature dimensions, each of which requires a separate learning channel. More theoretical work is needed to explore the scalability of the proposed model to settings with naturalistic inputs using vector-valued sensory prediction errors similar to those used in reinforcement learning (Lee et al., 2024), building on the simulations in Figure 9. Our emphasis on dopamine as a prediction error signal for learning does not preclude complementary roles that stem from sources of heterogeneity not considered here. Previous studies in the reward domain have proposed separate roles for both D1- and D2-expressing striatal neurons (Collins & Frank, 2014) and for tonic and phasic components of dopamine (Niv et al., 2007) in learning and instantaneous choice behavior. This spatiotemporal heterogeneity has recently been shown to have interesting consequences for learning due to differential dopamine affinities of D1 and D2 receptors (Pinto & Uchida, 2023). Incorporating such features into the proposed account of perceptual learning will be crucial for understanding the specific cell-type-specific mechanisms by which excess striatal dopamine leads to hallucinations and for developing safer treatments. Furthermore, incorporating known alterations in GABAergic/glutamatergic function within the proposed circuit model presents an opportunity to develop a unified computational theory of psychosis that integrates the dopamine hypotheses with the glutamatergic excitation/inhibition imbalance suggested by predictive processing models (Keller & Sterzer, 2024).

## 5 Methods

### 5.1 Model

We developed a model of parallel corticostriatal loops for learning sensory and reward statistics via a local plasticity rule in corticostriatal synapses, modulated by heterogeneous dopamine signaling in the striatum.

#### Dynamics

The model comprised two parallel corticostriatal loops – sensory and limbic – to enable decision-making by respectively learning sensory and reward statistics. The sensory loop included the sensory cortex and the sensory portion of the dorsal striatum i.e., the sensory striatum (SS), while the limbic loop included the frontal cortex and the ventral striatum (VS). The sensory cortex was modeled as a recurrent neural network with *N* units which combined time-varying external sensory inputs with the output of the sensory striatum to track the momentary belief state i.e., the posterior probability of the underlying state of the world given the observations, while *M* units in the sensory striatum are themselves driven by those in the sensory cortex. The frontal cortex was also modeled as a recurrent neural network which combined the belief state with the output of the ventral striatum to output an action that maximized reward. Analogous to the sensory striatum, *M* ventral striatal units are driven by *N* units in the frontal cortex. The dynamics of units in cortical and striatal regions in both loops are therefore given by the same set of equations where subscript *k* is used to index the corticostriatal loop (sensory or limbic):

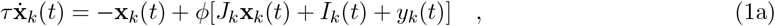

where *t* denotes time, *τ* is the intrinsic time-constant of each unit, nonlinearity *ϕ*(·) is the tanh function, **x**_*k*_ ∈ ℝ ^*N*^ denotes the population activity in cortical region indexed by *k, J*_*k*_ is the recurrent connectivity in that region, *I*_*k*_ is the scalar external input (sensory observations or posterior belief depending on the cortical region), and *y*_*k*_(*t*) = *σ*[**1**^*T*^ **z**_*k*_(*t*)] denotes the summed output of the striatal units in region *k* passed through a sigmoidal nonlinearity. Activity of striatal units **z**_*k*_ ∈ ℝ ^*M*^ is given by:

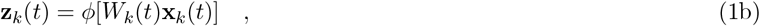

where *W*_*k*_ ∈ ℝ ^*M ×N*^ denotes the corticostriatal synaptic weights. Neural signatures of evidence accumulation for perceptual and value computations have been found in sensory association and frontal cortical regions respectively (de Lafuente et al., 2015; Lin et al., 2020). Accordingly, recurrent connectivity *J* within the cortical regions are pre-trained to perform perceptual inference i.e., belief computation (sensory cortex) or value-based decision-making i.e., action selection (frontal cortex) via backpropagation-through-time and then held fixed after this initial pre-training. Con-cretely, recurrent weights within the sensory cortex are pre-trained such that the summed activity encodes the posterior belief that the stimulus contains a signal i.e., to minimize the loss ∫{**1**^*T*^ **x**_*k*_(*t*) − *b*(*t*)}^2^ d*t* where *b*(*t*) = *P* (*s* = 1|*I*(1), …, *I*(*t*)) and *s* denotes the latent state (*s* = 1 for signal and *s* = 0 for noise). In contrast, recurrent weights within the frontal cortex are pre-trained such that the summed activity encodes the binary choice (yes/no) that maximizes reward i.e., to minimize the loss ∫{**1**^*T*^ **x**_*k*_(*t*) −*a*(*t*)}^2^ d*t* where *a*(*t*) = arg max_*a*_ *P* (*r* = 1|*b*(*t*_*d*_), *a*) ∀ *t > t*_*d*_ where *t*_*d*_ is the decision time. Critically, optimal performance depends on accurate estimation of sensory and reward expectations provided by the striatal output to the cortex i.e., *y*_*k*_(), which need to be continually learned in non-stationary environments. To enable this, the elements of corticostriatal connectivity matrix *W* in both loops are initially random (gaussian, sampled from *𝒩* (0, 1*/N*)) but modified over time by local plasticity as described below. Since the objective is to model the effect of striatal learning on cortical computation, we are agnostic to the precise temporal structure of individual striatal units and thus ignore recurrent inhibition within the striatum.

#### Plasticity rule

We assume a standard three-factor learning rule (Łukasz Kuśmierz et al., 2017; Gerstner et al., 2018) in which teaching signals encoded by dopamine serve as a third factor to gate plasticity in corticostriatal synapses according to:

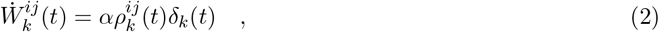

where 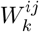 denotes the synaptic weight from cortical unit *j* to striatal unit 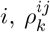 denotes the eligibility trace of that synaptic weight estimated as the covariance between the pre-synaptic and the (derivate of) post-synaptic activity: 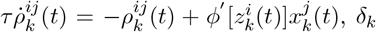 serves as the teaching signal (the third factor) to gate plasticity at that synapse, and *α* denotes the learning rate. Note that the prediction error signals are global in that they are independent of the identity of pre- and post-synaptic neuron (indexed by superscripts *i* and *j*). However, they are assumed to be region-specific (indexed by the subscript *k*) in keeping with the observed heterogeneity in dopamine signaling across striatal regions.

#### Optimization

Standard neural network approaches optimize network parameters (e.g., synaptic weights *W*) to maximize accuracy. Because our objective here is to develop a model that can adapt to changes in task statistics (i.e., stimulus and reward statistics), we instead optimize the teaching signal *δ*_*k*_ such that the local plasticity rule in Equation 2 above can enable the network to learn either the stimulus or the reward statistics. Because we optimize the model for learning rather than performance, this approach corresponds to learning-to-learn or metalearning. Concretely, we expressed the teaching signal as a spatiotemporal transformation of the activity of all units in the network:

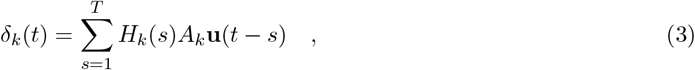

where **u**_*t*_ = (**x**_*t*_, **z**_*t*_)^*T*^ denotes the concatenated *P*-dimensional activity of all units in the network at time *t*, where *P* = 2*N* + 2*M, A*_*k*_ ∈ ℝ ^*K×P*^ and *H*_*k*_ ∈ ℝ ^*T ×K*^ respectively denote a rank-*K* spatial filter and temporal filter where *T* is the length of the filter (number of timesteps). *H*_*k*_(*s*) denotes row *s* of *H*_*k*_. To determine the teaching signal for learning sensory expectations, we optimized the elements of *A* and *H* using gradient descent in a signal detection paradigm in which the fraction of signal trials was resampled from a standard uniform distribution *U* (0, 1) every 20 trials for 10 blocks i.e., a total of 200 trials. To determine the teaching signal for learning reward expectations, we similarly optimized *A* and *H* but in a setting in which the fraction of trials with rewarded ‘yes’ responses was resampled from a standard uniform distribution *U* (0, 1) every 20 trials. In both cases, optimization (i.e., meta-learning) via gradient-descent was performed in an outer loop while the inner learning loop was used to update corticostriatal synapses according to Equation 2. We found that a single dimension (*K* = 1) was sufficient to optimize learning of sensory expectations while learning reward expectations required a two dimensional subspace (*K* = 2), and the simulations in the main text correspond to these choices. The composition of the optimized teaching signal was characterized by estimating the alignment between the columns of *A*_*k*_ and the directions of neural activity **u** encoding the belief (*b*), value (*v*), reward (*r*), and action (*a*).

#### Teaching signals

In addition to using the teaching signals derived via the optimization approach described above, we simulated the model using idealized teaching signals for learning sensory and reward expectations. In these simulations, we used sensory prediction error (sPE) and reward prediction error (rPE) as teaching signals for learning sensory expectation and reward expectation in the SS and VS respectively. sPE was defined as:

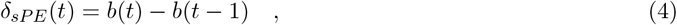

where *b*(*t*) = **1**^*T*^ **x**_*k*_(*t*) denotes the posterior belief encoded in the summed activity of the model sensory cortex as described earlier. Note that the teaching signal defined in this manner corresponds to the derivative of the subjective belief. rPE was defined according to the standard temporal difference learning rule:

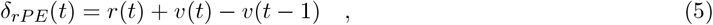

where *v*(*t*) denotes the expected reward associated with the current belief state and *r*(*t*) denotes the binary reward feedback.

#### Simulations

The cortical modules each comprise *N* =128 units while the striatal modules comprise *M* =16 units in keeping with convergent corticostriatal projections in the mammalian brain (Kincaid, Zheng, & Wilson, 1998). Simulations were performed using timesteps dt = 10ms, the intrinsic time constant of individual units was set to *τ* = 50ms (5 timesteps), and learning rate *α* = 0.1.

For simulating the associative learning task, the odor and reward were both modeled as brief 100ms pulses, spaced 1s apart. Since reward delivery coincided with the click of a water valve (Menegas et al., 2018), the sensory cortex received two sensory inputs in sequence. To qualitatively match the empirical findings, we initialized the sensory expectation for the water-valve click to be slightly higher than the odor, resulting in weaker sensory prediction errors. This assumption is justified since animals are likely to be more familiar with the former due to prior exposure to the experimental setup.

For simulating the signal detection task, the stimuli were 500ms long and comprised 10 samples of 50ms each. Samples were drawn from a gaussian with mean *µ* ∈ {0, 0.1} and variance *σ* ∈ {0.1, 0.2, 0.3} where mean and variance controlled the trial type (signal or noise) and signal to noise ratio respectively. For simulations used to test the model’s performance on signal detection, we used two blocks of 500 trials each. In the first block, the fraction of signal trials was 0.5 and rewards were delivered with a probability of 0.5 both for correct ‘yes’ (true positive) and correct ‘no’ (true negative) responses. In the subsequent block, we either increased the fraction of signal trials to 0.75 (signal manipulation) or the fraction of rewarded ‘yes’ trials to 0.75 (reward manipulation).

To simulate excess dopamine, we introduced multiplicative perturbation to the teaching signals as 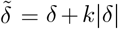, where the positive constant *k* determines the strength of the perturbation (Maia & Frank, 2017). Note that this form of perturbation can be re-written as:

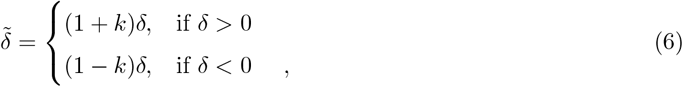

implying that this is equivalent to introducing asymmetric learning rates where trials with positive prediction errors have a stronger influence on learning. Learning rate asymmetry for different stimuli and/or outcomes is consistent with studies on the impact of pharmacological pro-dopaminergic drugs on learning and decision-making (Rutledge et al., 2009; Pagnier et al., 2024). We used *k* = 0.45 and *k* = 0.35 to recapitulate the effect of CAPS on human behavior in the signal detection task and conditioned hallucinations task (Powers et al., 2017) respectively.

For simulating the ten-way classification task, we first pre-trained an RNN model to output the momentary joint posterior distribution over the classes given the time course of ten-dimensional sensory observations where the target distribution was defined in accordance with Bayes’ rule, and a ten-dimensional sensory expectation. Each feature dimension of the observation was provided via a separate input channel such that all but one channel had a non-zero mean for observations from a given class. Likewise, different dimensions of sensory expectation were provided as inputs from separate striatal units. After pre-training with data drawn from a uniform distribution over classes, we tested the model by modifying the input distribution to be a spike-and-slab distrbution where observations in half the trials were blank i.e., all channels had zero-mean inputs, while the other half were drawn from an exponential distribution over classes. The plasticity rule in corticostriatal synapses was given by equation 2.

### 5.2 Mouse Behavioral Tasks

#### Associative Learning

As described in greater detail in the original work (Menegas et al., 2017), bulk calcium signals were recorded from dopamine axons projecting to the ventral striatum or tail striatum while mice learned associations between an odor and an outcome. Importantly, the mice had no prior experience with the odor. Each trial, an odor was presented followed by a one second delay and then either a water reward (90% of trials) or nothing (10% of trials).

#### Auditory Signal Detection

As described in greater detail in the original work (Schmack et al., 2021), on each trial water-restricted mice reported their auditory percept (either a 1 kHz tone or white noise) by pressing a lever. Tone trials were presented on 50% of trials. Correct decisions were rewarded with water after a random delay period. Trials were self-initiated, which created tension between waiting for a reward and initiating another trial. This willingness to wait for reward has been previously validated as a measure of statistical decision confidence (Hangya et al., 2016).

### 5.3 Open-Source Human Tasks

#### Random Dot Motion Task

As described in greater detail in the original work (Zylberberg et al., 2018), on each trial participants reported whether a cloud of dots were moving leftward or rightward and their confidence. The duration of dot presentation and motion coherence were varied across trials. Within each block (15-42 trials), one motion direction was presented more often. Participants were not provided feedback about their performance or the block identity until the end of each block.

#### Conditioned Hallucinations Task

As described in greater detail in the original work (Powers et al., 2017), on each trial participants were presented with a visual checkerboard stimulus and a coincident 1 kHz pure tone in white noise. In the beginning of the task, the visual checkerboard reliably predicted the presence of the tone, but as the task progressed, the probability of tone presentation was gradually decreased. Participants reported whether the tone was present and their confidence. Participants did not receive feedback about their performance.

### 5.4 Human Data Collection

#### Sample Information

All participants provided informed consent and all study procedures were approved by the local Institutional Review Board. For both samples, participants were recruited through Prolific and the experiment was hosted on Pavlovia.org (Peirce et al., 2019).

#### Participant Pre-screening and Retained Sample

The distribution of subclinical hallucination propensity is heavily skewed in the general population, so participants with high hallucination propensity are poorly represented in random samples. To ensure a high between-subject variability of hallucination propensity, and in line with best sampling practices (Liu, Abdellaoui, Verweij, & van Wingen, 2023; Kang et al., 2024), we used a pre-screening procedure following our previous work (Ashinoff, Buck, Woodford, & Horga, 2022). We first collected large initial samples (N=499 and N=494 for Sample 1 and Sample 2, respectively) which completed the Cardiff Anomalous Perception Scale (CAPS) and the Beck Depression Inventory (BDI, which we used to approximately match groups on general psychopathology). Then, using previous published norms (Schmack et al., 2021) we derived unbiased CAPS-score cutoffs to determine three groups: low, medium, and high (under 30.1 for low and over 85.7 for high). All participants from the high CAPS group and matched participants in the low and medium CAPS groups (according to their BDI score) were invited to complete the behavioral task. 152 and 148 participants completed the full experiment in Sample 1 and Sample 2, respectively.

#### Human Auditory Signal Detection

To maximize comparisons with the mouse data, core features of the human task were matched to the mouse task. White noise was constantly played in the background and on some trials, a pure 1 kHz tone was presented. Detection difficulty was standardized across participants by using a separate auditory thresholding procedure (Powers et al., 2017) to determine the volume at which the tone was detected on 25%, 50% and 75% of presentations. To further standardize the delivery of auditory stimuli, participants were required to wear headphones which was enforced using a headphone check procedure based on prior work (Huggins Pitch and Binaural Beats, (Milne et al., 2021)).

To evaluate the influence of sensory and reward learning on high-confidence false alarms, two blockwise manipulations were introduced in the human task (**Figure 5A** – right): 1) the probability of a tone being presented (i.e., signal probability), and 2) the probability of receiving additional reward for a signal decision (i.e., reward probability). Since using these contingencies maximized overall reward receipt, participants were incentivized to engage in both sensory and reward learning and incorporate their learned expectations into their choices and confidence reports. Importantly, we selected block probabilities to minimize the correlation between perceptual correctness and reward magnitude which allowed us to infer distinct effects of both signal and reward probability on behavior. The task consisted of 12 blocks of 20 trials each. There were three categories of blocks: signal, reward, and hybrid manipulation blocks. In signal blocks, reward probability is fixed at 50% and signal probability is either 10%, 30%, 70%, or 90% (4 blocks total). In reward blocks, signal probability is fixed at 50% and reward probability either 10%, 30%, 70%, or 90% (4 blocks total). In hybrid blocks, signal and reward probability are either 20% or 80% (all combinations result in 4 blocks total).

To further separate the influence of sensory and reward contributions to choice, participants also reported how certain they were that a tone was present (i.e., signal confidence) on a visual analog scale. At the end of each trial, participants were told whether the tone was indeed present or absent and whether their choice resulted in an additional reward (**Figure 5A** – left).

#### Questionnaires and Additional Cognitive Tasks

In the same session as the auditory signal detection task, participants repeated the CAPS and BDI and also completed the Beck Anxiety Inventory, Raven’s progressive matrices, and the forward and backward digit span. We used the Raven’s score to test for specificity of hallucination-proneness effects.

#### Data Quality

To ensure task comprehension, participants were required to complete several practice sessions and a quiz before completing the signal detection task. For participants who completed the full task, we also used self-report and task-based screening measures to promote high data quality (Zorowitz, Solis, Niv, & Bennett, 2023). Each self-report questionnaire included a catch question which should elicit a standard response from attentive participants. Participants were excluded if two or more responses to catch questions were incorrect (N=1 and N=1 for Sample 1 and Sample 2, respectively). To promote the reliability of hallucination propensity measures, participants were also excluded for improbably large changes from the first to second administration of CAPS (changes in group assignment based on normed cutoffs from low to high or high to low; N=2 and N=1 for Sample 1 and Sample 2, respectively). Finally, participants were excluded for suspicious median response times (*<*200 ms or *>*3 s) during the signal detection task (N=4 and N=2 for Sample 1 and Sample 2, respectively). With these exclusions, the final sample sizes used for analyses were N=146 and N=144 for Sample 1 and Sample 2, respectively.

### 5.5 Data Analysis

In general, we use mixed-effects regression models to evaluate the influence of variables of interest on behavior or neural data at the group-level: *Y*_*i,j*_ = *β*_0_ + *β*_1_*X*_*ij*_ + *u*_0*j*_ + *ϵ*_*ij*_ where the outcome *Y* for subject *j* on trial *i* is the determined by the predictor *X*_*ij*_, a group-level intercept *β*_0_, a group level slope *β*_1_, and subject-specific random intercepts *u*_0*j*_. When modeling choice, we use mixed-effects logistic regression, while for confidence and neural activity, we use mixed-effects linear regression. All continuous variables were z-scored prior to modeling. To limit Type 1 errors, coefficient-level confidence intervals and p-values were estimated using parametric bootstrapping, with *B* = 2000 simulations per model (Yu et al., 2022). p-values reflect the proportion of bootstrap estimates with the opposite sign from the observed effect (two-sided). Since the resolution of this test is limited by the number of simulations, effects with no sign reversals are reported as *p <* 1*/*(*B* + 1) (i.e., *p <* 5 *×* 10^−4^ for 2000 simulations).

#### Mouse Fiber Photometry Analysis (Data from Schmack et al., 2021)

A prediction error should be composed of a positive response to a stimulus/outcome and a dissociable negative response that scales with the expectation of the stimulus/outcome (Rutledge, Dean, Caplin, & Glimcher, 2010). So, to test whether stimulus-evoked dopamine release in the striatum was consistent with a sensory prediction error, we evaluated whether changes in these signals during stimulus presentation were sensitive to stimulus history (an index of sensory expectation) after adjusting for stimulus signal-to-noise ratio, reward history, and choice history. Specifically, we tested whether dopamine signal during a region-specific window starting at stimulus onset was modulated by SNR, previous trial stimulus, previous trial reward, and previous trial choice. To reduce cross-trial variability, dopamine signal from a 100 ms pre-stimulus baseline was subtracted from the signal in the region-specific window. The region-specific window length was defined as the half-width of signal autocorrelation estimated at the group level. For exploratory visualization, we used moving window regressions with 500 ms windows incremented by 250 ms. Dopamine signals were normalized as Δ*F/F*, computed as the relative fluorescence change from a local baseline (median-filtered raw signal over a 60s window). See (Schmack et al., 2021) for details.

#### Optogenetic stimulation Analysis (Data from Schmack et al., 2021)

(Schmack et al., 2021) reported that optogenetic stimulation of dopamine release in the tail striatum increased false alarm rate and confidence. To assess more granular temporal dynamics of this effect, we predicted choice and confidence as a function of experiment progress (i.e., session number), session progress (i.e., block number), and block progress (i.e., trial number within a block). Since the number of sessions varied across animals and the number of blocks varied across animals and sessions, we normalized predictors by taking median splits for each animal. In other words, we considered whether the effect of dopamine stimulation changed from the first to the second half of sessions, blocks, or trials.

#### Self-Supervision Analysis (Data from Powers et al., 2017 and Zylberberg et al., 2018)

Under our model, sensory expectations are updated in a self-supervised manner based on trial-by-trial changes in belief. To evaluate empirical evidence for this assumption, we tested how belief history influenced current perception in perceptual tasks without explicit feedback. Specifically, the model predicts that choices are more likely to be repeated when previous confidence was high, regardless of the current trial stimulus strength (e.g., signal-to-noise ratio or coherence) and task context (e.g., base rate). Accordingly, we evaluated whether the probability of repeating a choice was related to previous trial confidence (binned for each subject).

#### Human Conditioned Hallucinations Task Analysis (Data from Powers et al., 2017

To connect our model to standard algorithmic modeling approaches in the field, we also fit a sequential learning model to choice data. Under this model, a perceptual posterior is computed as a weighted combination of current sensory information (likelihood) and prior expectations. The likelihood function was defined as Gaussian distribution with mean *µ* and standard deviation *σ*. The prior corresponded to a point estimate of the relative presence of signal. The relative weighting of priors versus likelihood was parameterized by a tradeoff parameter *ω*. All three parameters (*µ, α, ω*) were fit to each subject’s choice data.

#### Human Auditory Signal Detection Analysis

Participants are incentivized to learn the sensory and reward contingencies of the task to develop expectations about the sensory environment and which decisions yield additional reward. If participants are incorporating their learned expectations into their behavior, recent sensory or reward evidence should measurably bias choice and confidence. To quantify this effect, we predicted current trial choice and confidence as a function of SNR, previous trial sensory feedback, previous trial reward outcome. To adjust for non-learning effects like choice perseveration, we also tested models that include previous trial choice as a predictor. To evaluate whether learning effects are moderated by hallucination proneness, we also tested models that included interactions of each predictor with total scores on the Cardiff Anomalous Perceptions Scale (CAPS) and/or the Raven’s progressive matrices into the mixed-effects regression models.

Choices are influenced by perceptual beliefs and the mapping between these beliefs and actions (i.e., policy). Since confidence reports are a readout of perceptual beliefs we can estimate participants’ policy by evaluating how our reward manipulation influences choice after accounting for the effect of confidence. To this end, we predicted choice as a function of SNR, previous trial sensory feedback, previous trial reward outcome, and current trial confidence. Since confidence is a separate predictor in this model, we interpret any influence of reward history on choice as an effect of a change in policy.

## 6 Acknowledgements

We thank Katharina Schmack and Adam Kepecs for giving access to their data published in (Schmack et al., 2021), and Albert Powers for sharing their data published in (Powers et al., 2017). We thank Arturo Torres for helpful discussions. This work was supported by NIH grants R01MH136672 to G.H. and C.K. and F31MH134617 to J.B., the Kavli Foundation, the Gatsby Charitable Foundation GAT3780, and a NARSAD Young Investigator Grant #31985 from the Brain & Behavior Research Foundation to K.L.

