## Supplemental Figures for "A corticostriatal learning mechanism linking excess striatal dopamine and auditory hallucinations"

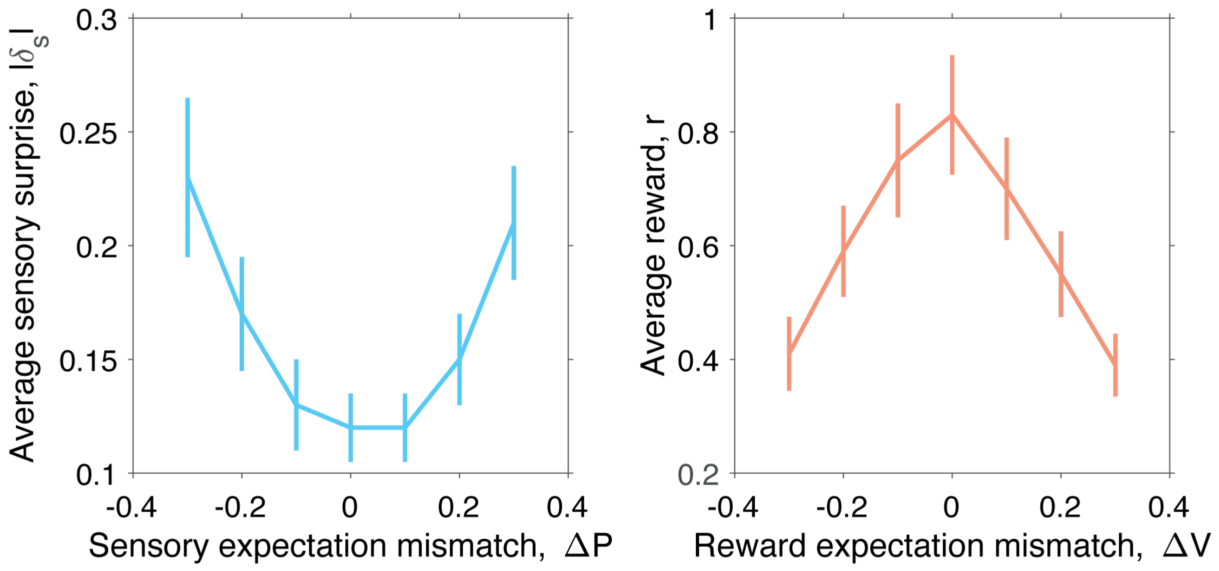


**Fig S1. Left**: The average magnitude of sensory surprise (per trial) incurred by the model as a function of the degree of mismatch between actual and assumed sensory expectation (P). **Right**: The average amount of reward (per trial) gained by the model as a function of the degree of mismatch between actual and assumed reward expectation (V). Positive and negative mismatch correspond to optimistic and pessimistic expectations respectively. Error bars denote standard error in mean estimated by bootstrapping.

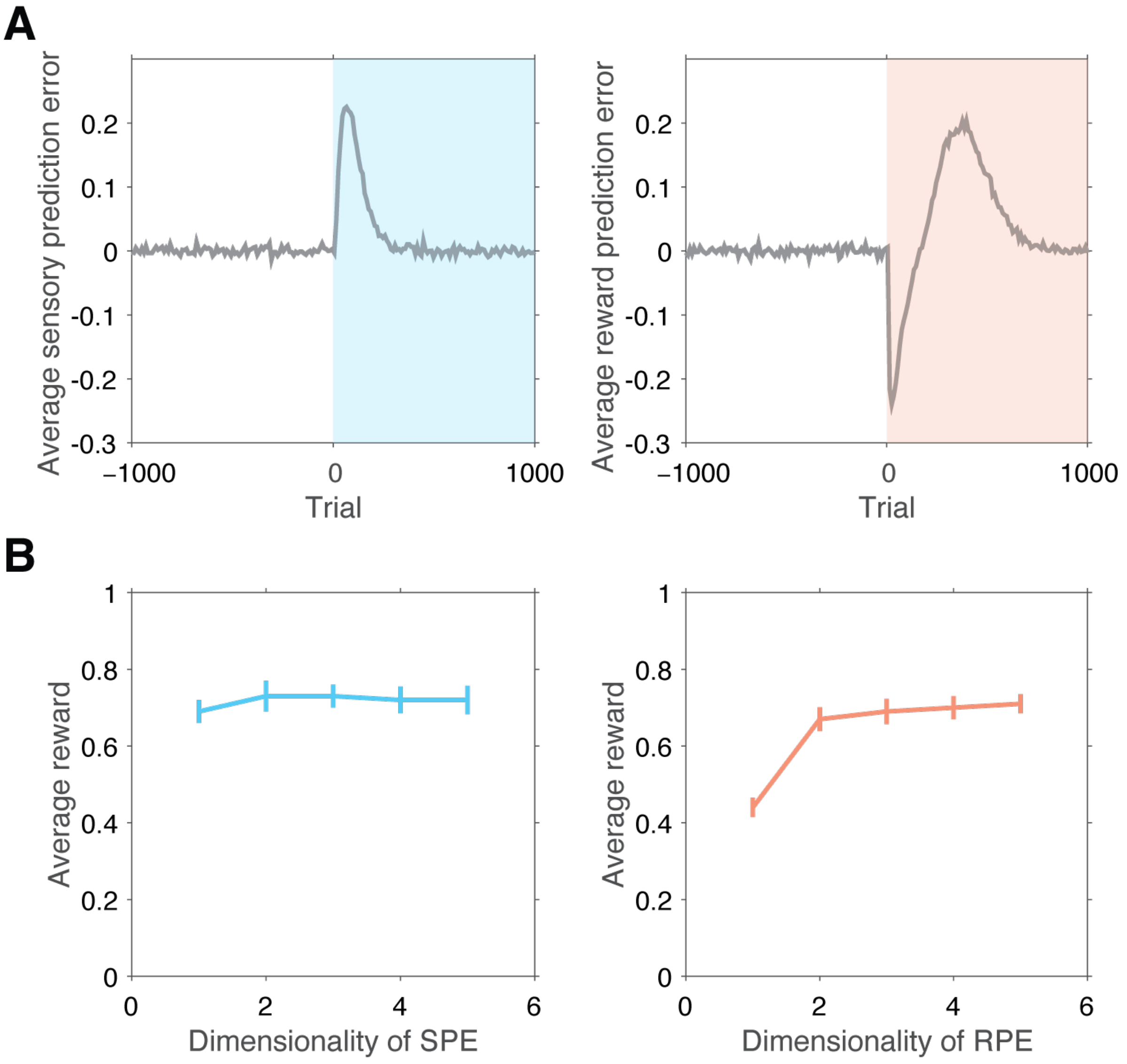


**Fig S2. (A)**. **Left**: Average sensory prediction error as a function of trials since reversal of signal statistics (i.e., proportion of signal trials). **Right**: Average reward prediction error as a function of trials since reversal of reward statistics (i.e., relative proportion of 'yes' vs 'no' responses that are rewarded). **(B)** Average reward gained by the model as a function of the dimensionality of the optimized sensory prediction error (**left**) and the dimensionality of the optimized reward prediction error (**right**) respectively. The dimensionality was controlled by varying the number of spatiotemporal filters (*K*) that are optimized by the meta-learning procedure (methods).

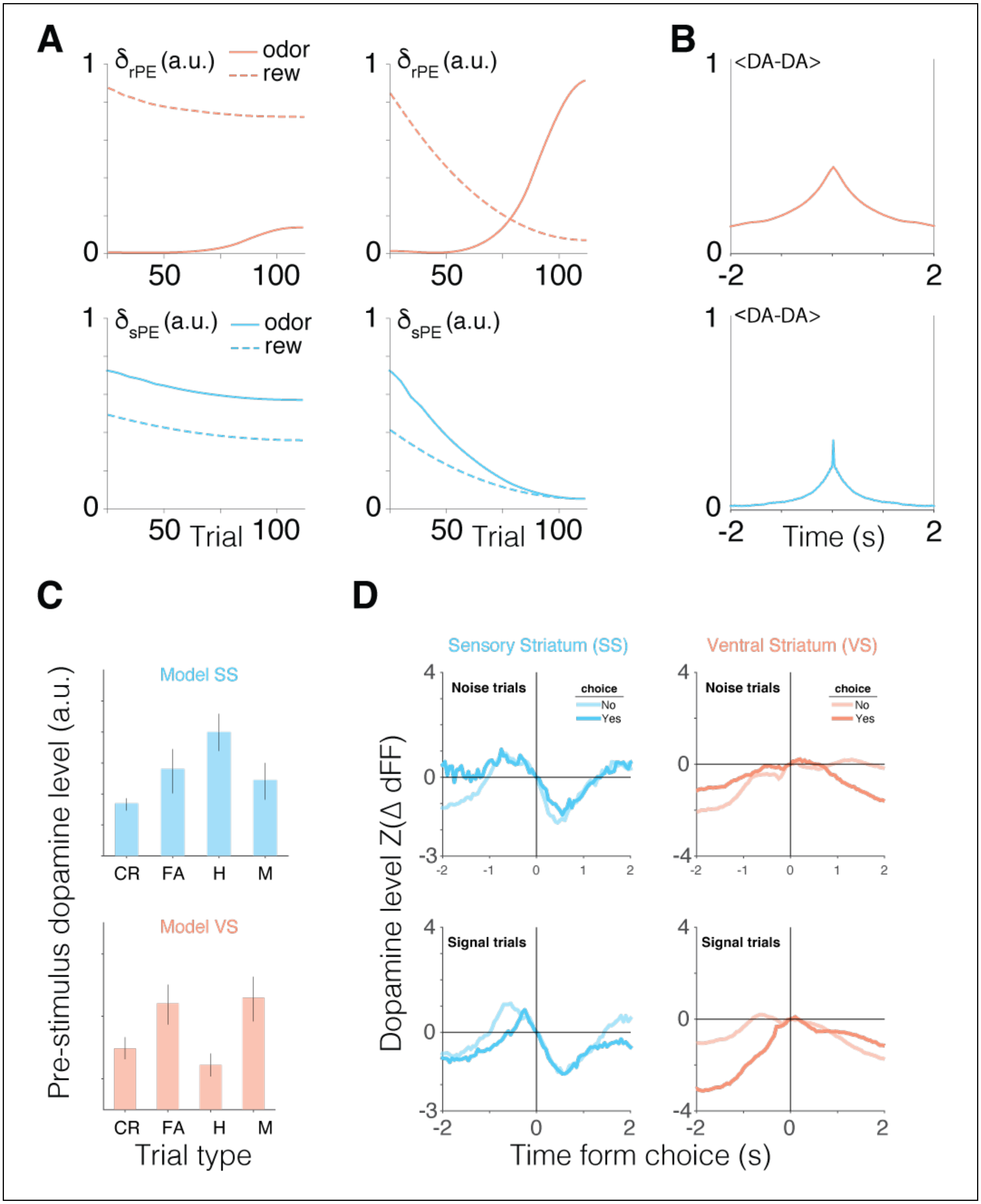
A
δrPE (a.u.)
odor
rew
1
0
50
100
B
<DA-DA>
-2
2
δsPE (a.u.)
Trial
Time (s)
C
Model SS
Model VS
Pre-stimulus dopamine level (a.u.)
CR
FA
H
M
Trial type
D
Sensory Striatum (SS)
Ventral Striatum (VS)
Noise trials
Signal trials
choice
No
Yes
Dopamine level Z(Δ dFF)
Time form choice (s)

**Fig S3. (A)**. **Left**: Evolution of the amplitude of dopamine transients evoked by odor (solid lines) and reward (dashed lines) in the model ventral striatum (top) and sensory striatum (bottom) respectively with the discount factor set to $\gamma = 0.05$. **Right**: Similar to the left panels, but for simulations with the discount factor set to $\gamma = 0.95$. **(B)**. The auto-correlogram of the dopamine responses in the ventral striatum (top) and the tail of the striatum (bottom), averaged across sessions. The full-width at half-maximum was used as define the length of the time window for estimating the time-averaged magnitude of stimulus-evoked dopamine transients in these regions. **(C)** Average pre-stimulus dopamine levels in the model SS (top) and VS (bottom) across trials grouped by stimulus and choice (CR: Correct Rejection, FA: False Alarm, H: Hit, M: Miss). Zero-mean additive noise was added to the model cortical neural activity in these simulations. **(D)**. Dopamine signals in the two regions averaged across trials grouped by stimulus (signal or noise) and choice (yes or no), aligned to the time of choice. Action prediction error should manifest as transients at the time of choice.

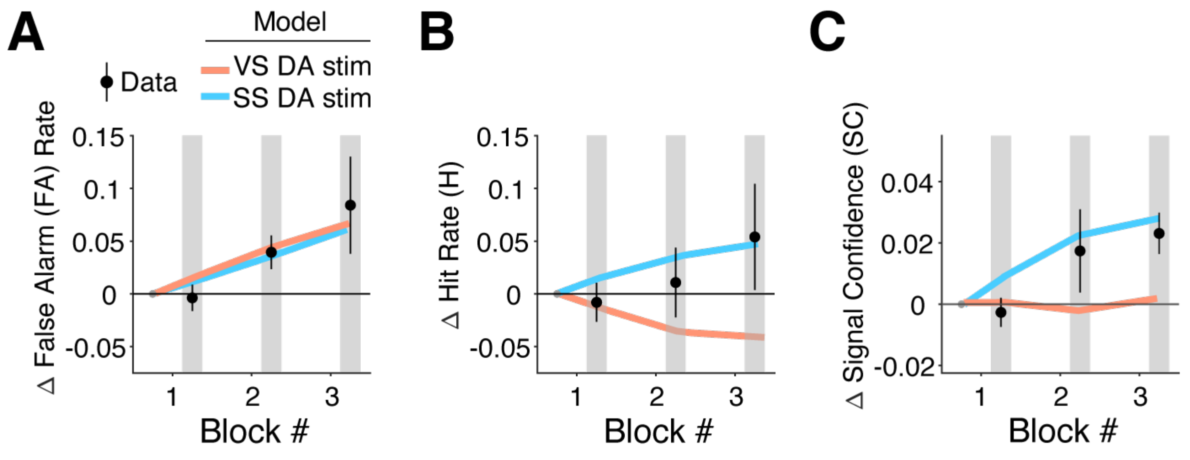


**Fig S4.** Model simulations with multiplicative perturbation of dopamine signals in the ventral striatum (VS) and the sensory striatum (SS), overlaid on mouse data. **(A)**. Change in the rate of false alarms with respect to the baseline (i.e., the first block of the experimental session) in each block. **(B)**. Similar to A but showing the change in the rate of hits. **(C)**. Similar to A but showing the change in signal confidence.

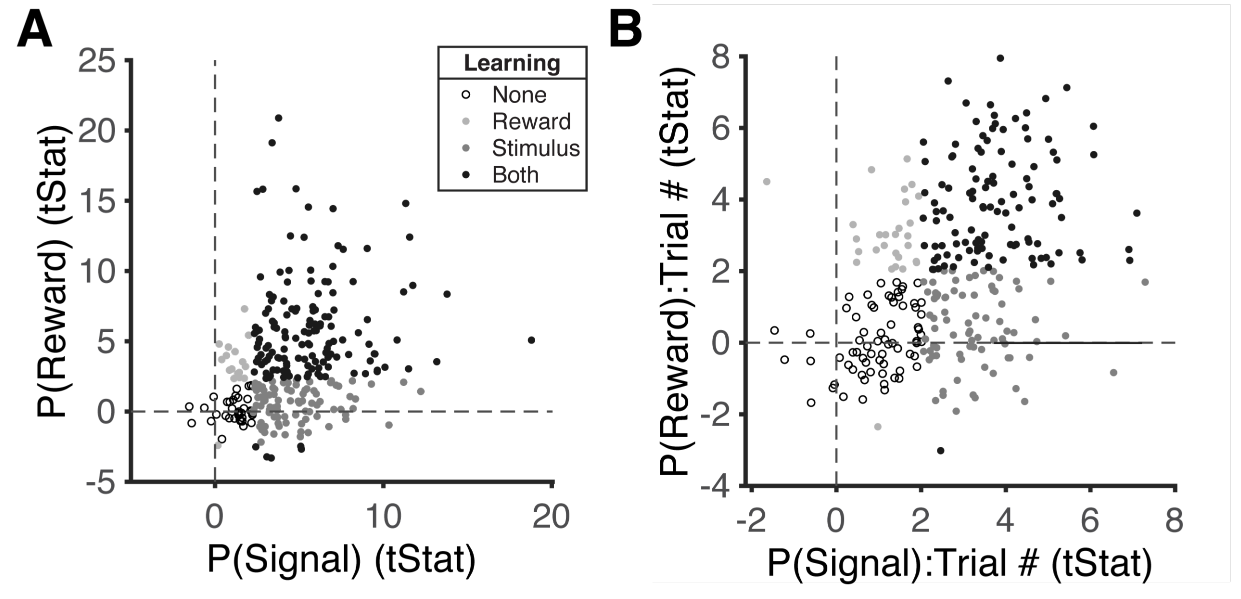


**Fig S5. (A)**. The strength of the regression capturing the influence of the true block-level signal and reward probabilities on each participant's subjective rating at the end of the block, reflecting the asymptotic learning performance of participants. **(B)**. Similar to (A) but showing the strength of the interaction between subjective rating and trial number, reflecting the gradual learning process of participants. *Open circles – participants with no significant influence of either statistic; Light gray – significant influence of reward but not signal probability; Dark gray – significant influence of signal but not reward probability; Black – significant influence of both signal and reward probabilities.*

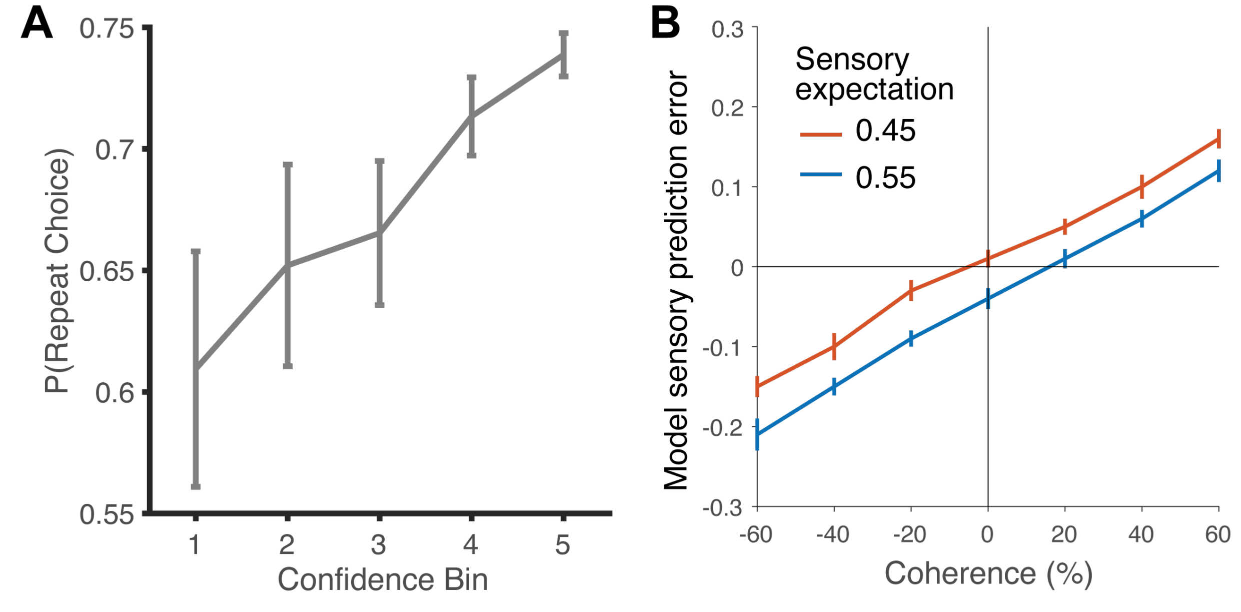


**Fig S6.** Motion discrimination task. **(A)** The probability of repeating the previous trial's choice (left/right) as a function of the confidence level in the previous trial discretized into five bins. Error bars denote standard error of the mean across participants. **(B)** Model sensory prediction error for trials grouped by coherence under two different levels of sensory expectation (a priori belief in positive coherence stimulus). Uninformative stimuli (0% coherence) increases or decreases the sensory expectation when a priori expectation is below or above 0.5 respectively.

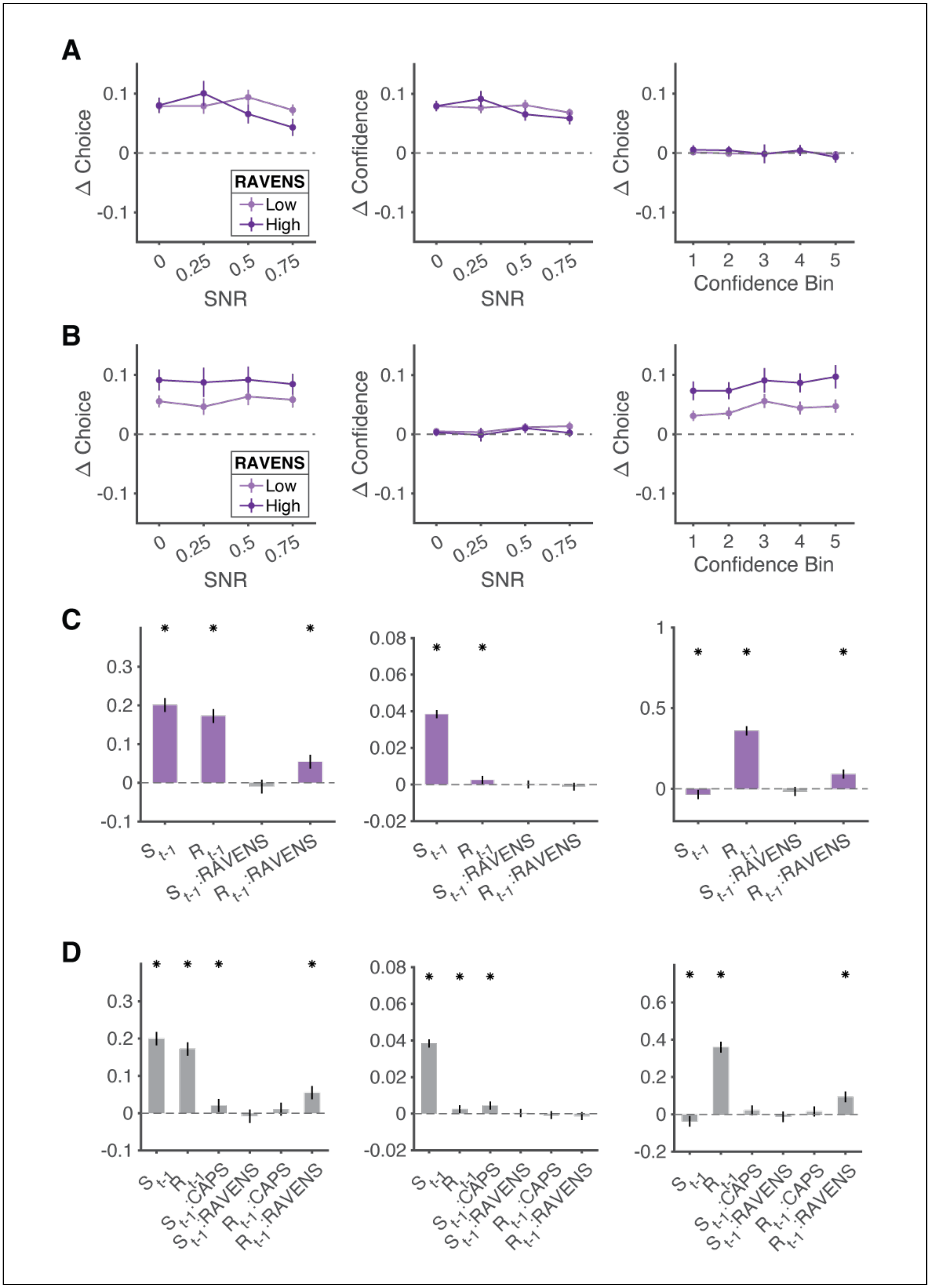

A
B
C
D
Δ Choice
Δ Confidence
RAVENS
Low
High
SNR
Confidence Bin
S t-1
R t-1
S t-1:RAVENS
R t-1:RAVENS
S t-1:CAPS
R t-1:CAPS

**Fig S7.** Cognitive deficits exaggerate reward-history effects. **(A)**. Left: The change in the rate at which participants report hearing a signal as a function of signal to noise ratio, after a signal trial, relative to after a noise trial. Middle: Similar to the left panel, but showing the change in the reported perceptual confidence. Right: Similar to the left panel, but showing the change in policy i.e., rate at which participants report hearing a signal as a function of confidence. Participants are grouped by their RAVENS scores. **(B)**. Similar to (A) but showing the change in response measures due to the previous trial's reward feedback. **(C)**. Coefficients of a mixed effects model, showing the effect of previous trial's stimulus ($s_{t-1}$) and reward ($r_{t-1}$), cognitive deficit (RAVENS), and the interactions ($s_{t-1}$ × RAVENS, $r_{t-1}$ × RAVENS) on choice (left), confidence (middle) and policy (right). **(D)**. Similar to (C) but showing the coefficients of a mixed effects model that additionally included the interactions of previous trial's stimulus and reward with hallucination proneness (CAPS). Asterisks denote $p < 0.05$. For A and B, error bars denote standard error of the mean across participants and for C and D, error bars denote 95% confidence intervals.

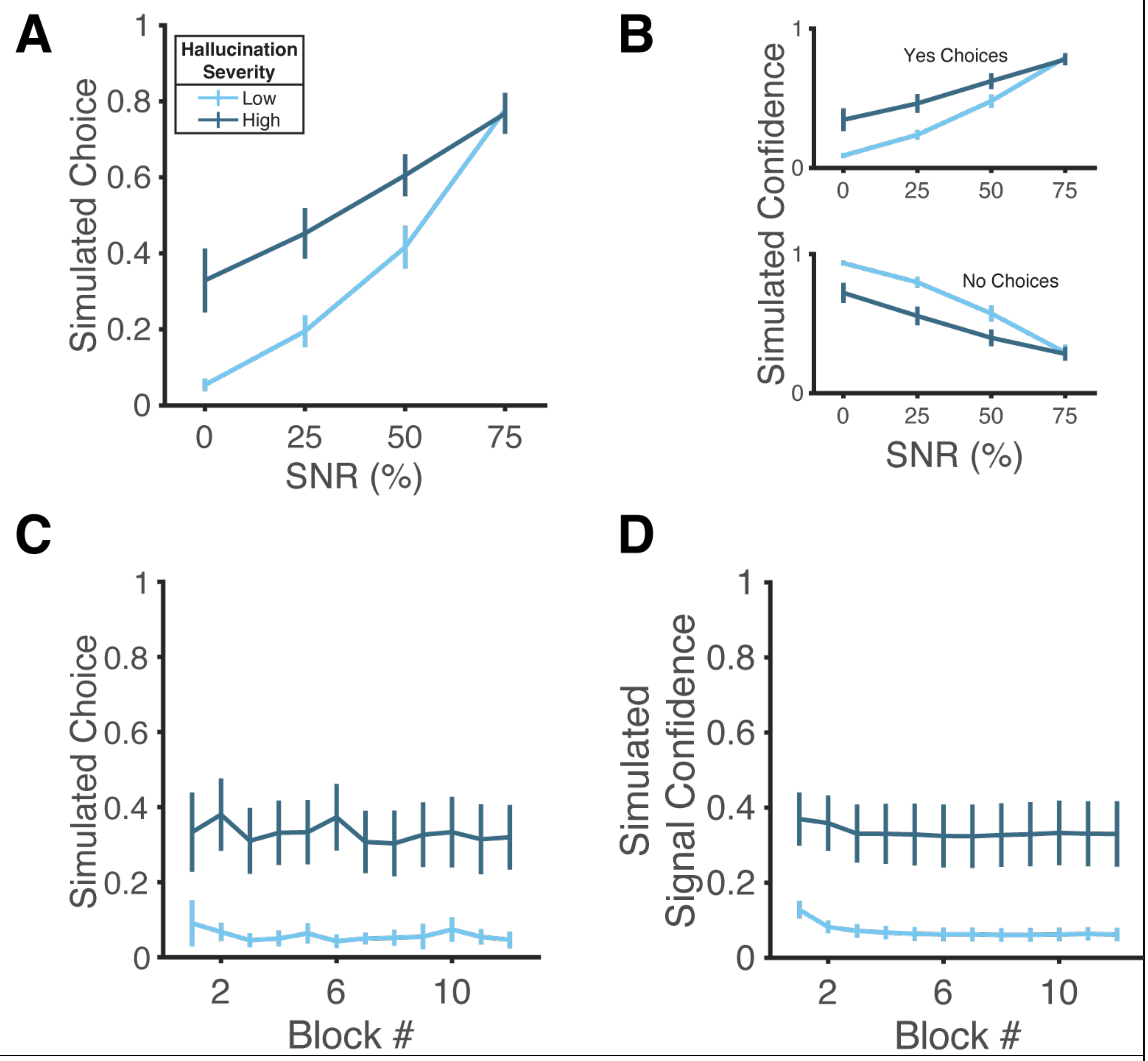


**Fig S8.** Simulations of the best-fit bayesian models. **(A)**. Proportion of 'yes' choices as a function SNR for patients grouped by hallucination severity (light vs dark, Methods). **(B)**. Model simulated confidence levels, shown separately for trials grouped by the type of response ('yes' or 'no'). **(C)**. Evolution of the proportion of false alarms across blocks. **(D)**. Evolution of the signal confidence across blocks. Error bars denote standard error of the mean across participants.
